# Brain pericytes are an oxidative stress-sensitive cell population damaged through ROS-dependent TRPA1 activation

**DOI:** 10.64898/2026.10.19.760199

**Authors:** Christopher J. Groten, Jasmine Kwan, Julia Groening, Maya Bellville, Nicholas L. Weilinger, Yanyang Bai, Stefan Wendt, Zoe Kortje, Faithful Olarinde, Isabelle Chung, Hyun B Choi, Louis-Philippe Bernier, Ravi L. Rungta, Brian A. MacVicar

## Abstract

Oxidative stress is a pathological process driven by an imbalance between the production and clearance of reactive oxygen species (ROS). It is a hallmark of numerous brain disorders, driving neuroinflammation, blood-brain barrier (BBB) disruption, and neurodegeneration. Here, we used two-photon microscopy in rodent brain slices to evaluate cell-type vulnerability to oxidative stress. We identified a microvascular cell population displaying pronounced sensitivity to H_2_O_2_ and thiol oxidation. These Pdgfr-β expressing cells were identified as pericytes and perivascular fibroblasts. Because pericytes regulate capillary perfusion and BBB integrity, we examined mechanisms driving their injury. Pericyte damage was independent of apoptosis or necroptosis, but required Ca^2+^ influx. While blocking voltage-gated Ca^2+^, TRPC, Orai channels was ineffective, inhibitors of TRPA1 channels reduced pericyte damage. Immunostaining and western blotting showed TRPA1 associated with brain capillaries and pericytes. Functionally, TRPA1 regulated uptake of the pericyte-selective marker NeuroTrace, whereas pericyte GCaMP6f imaging demonstrated that TRPA1 was not the primary driver of cytosolic Ca^2+^ elevations during oxidative stress. These findings establish pericytes as vulnerable targets of oxidative stress and implicate TRPA1 as a mediator of pericyte structural degeneration. This sensitivity to oxidative stress, mediated by ROS-TRPA1 coupling, could contribute to the pericyte stress and degeneration observed in brain pathology.

## Introduction

A wide range of neurological disorders exhibit evidence of oxidative stress in the brain ^1–3^. Oxidative stress is defined as an imbalance between the production and removal of reactive oxygen species (ROS), resulting in cellular dysfunction ^4^. ROS comprise both radical and non-radical oxygen-containing molecules, including superoxide, hydrogen peroxide, and hydroxyl radicals ^4^. In the brain, ROS are generated by multiple cell types through NADPH oxidases, metabolic enzymes, and the mitochondrial electron transport chain^5^. Excessive ROS induce oxidative modifications of proteins, lipids, and nucleic acids, leading to impaired cellular function and viability ^6^. These oxidative modifications have been documented in numerous neurological disorders, including Parkinson’s disease, Alzheimer’s disease, and amyotrophic lateral sclerosis (ALS)^1–3^. Furthermore, mutations in genes involved in redox homeostasis have been linked to various brain disorders^7–9^. Despite extensive evidence implicating oxidative stress in neurological disease, its causal contribution to disease progression remains unclear. Reducing oxidative stress has shown beneficial effects in animal models of stroke, ALS, and Alzheimer’s disease, yet antioxidant therapies have largely failed in human clinical trials for these disorders ^7,10,11^. These findings highlight the need for a better understanding of the cellular and molecular mechanisms by which ROS impair brain function, with the goal of identifying new therapeutic targets that mitigate the downstream consequences of oxidative stress.

While reactive oxygen species (ROS) serve essential physiological signaling roles, excessive ROS in the central nervous system trigger widespread cellular dysfunction and death. Several mechanisms mediate this toxicity, including plasma membrane disruption, mitochondrial dysfunction, and ionic dysregulation via redox-sensitive channels such as pannexins and transient receptor potential (TRP) channels ^12–16^. These pathological effects impact virtually every constituent of the brain parenchyma-disrupting neuronal networks, provoking inflammatory and degenerative responses across glial populations, and impairing cerebrovascular function through endothelial cells and pericytes. Although oxidative stress affects diverse cell types, evidence indicates that vulnerability across the parenchyma is non-uniform, with select cell populations exhibiting heightened sensitivity to oxidative damage ^17–19^. Systematically resolving which cell populations are most vulnerable and determining the molecular mechanisms driving that susceptibility is essential for understanding the pathogenesis of oxidative stress-associated brain disorders.

In this study, we aimed to characterize cell-specific damage induced by oxidative stress in brain tissue and identify mechanisms that protect against this injury. Using two-photon microscopy, we visualized cellular damage following acute exposure to oxidative stressors and identified a microvascular-associated cell population that was damaged by oxidative stress.

These cells expressed Pdgfr-β and were located predominantly on capillaries, corresponding to pericytes, with another cell population on primary order vessels and low branch order vessels corresponding to perivascular fibroblasts. Due to the abundance of capillaries and critical importance of pericytes in brain disease, we focused on identifying the mechanisms underlying oxidative damage to pericytes. We identify that pericyte injury in oxidative stress is driven by Ca^2+^ influx and is mediated by TRPA1 channels, highlighting a mechanism of vascular vulnerability and a potential therapeutic target for limiting oxidative damage in neurological disease.

## Materials and Methods

### Animals

Sprague Dawley rats were used for this study and were purchased from Charles River Laboratories. The mice strains used were C57BL/6J (Jackson Laboratory) and GCaMP6f; ATP13a5-CreERT2 mice. The ATP13a5-CreERT2 mice were a gift from Dr. Zhen Zhao (University of Southern California) and crossed with Ai95 (RCL-GCaMP6f)-D reporter mice (Jackson Laboratory) to obtain GCaMP6f; ATP13a5Cre; double transgenic mice, which express GCaMP6f in pericytes in a Cre-dependent manner. To induce GCaMP expression, mice received intraperitoneal (IP) injections of tamoxifen (100 mg/kg) once daily for five consecutive days.

Tamoxifen was dissolved in corn oil and administered at a volume of 10 mL/kg body weight. Rat and mouse housing and experimental procedures were performed in accordance with Canadian Council on Animal Care (CCAC) regulations and protocols approved by the University of British Columbia and University of Montreal Animal Care Committees. Animals were group housed, fed ad libitum, and kept on a 12/12 hr light/day cycle. Both male and female rats (aged P17-P50) and mice (P28-P163) were used for our experiments.

### Acute brain slice preparation and solutions

Animals were anesthetized with isoflurane in an induction chamber and subsequently decapitated. The brains were then quickly removed and placed in ice-cold slicing solution containing the following (in mM): 120 N-methyl-D-glucamine, 2.5 KCl, 25 NaHCO_3_, 1 CaCl_2_, 7 MgCl_2_, 1.25 NaH_2_PO_4_, 20 D-glucose, 2.4 sodium pyruvate, and 1.3 sodium L-ascorbate. The slicing solution was continuously oxygenated with 95% O_2_ and 5% CO_2_. Brains were then cut with a vibratome (Leica VT1200S) into coronal slices (300 µm thick) of the cortex. Brain slices were immediately transferred to a chamber containing artificial cerebral spinal fluid (aCSF), which was warmed to 32°C and continuously oxygenated with 95% O_2_ and 5% CO_2_. aCSF contained (in mM): 126 NaCl, 2.5 KCl, 26 NaHCO_3_, 2 CaCl_2_, 1.5 MgCl_2_, 1.25 NaH_2_PO_4_, and 10 D-glucose, pH 7.3–7.4, osmolarity 310 mOsm. A Ca^2+^-free aCSF solution was used to test the involvement of extracellular Ca^2+^ entry and contained (in mM): 126 NaCl, 2.5 KCl, 26 NaHCO_3_, 3.5 MgCl_2_, 1.25 NaH_2_PO_4_, 10 D-glucose, 1 EGTA.

For pericyte GCaMP imaging in mouse cortical slices, the same general preparation procedure was used with minor modifications to the slicing and recovery solutions. The NMDG-based slicing solution contained (in mM): 120 N-methyl-D-glucamine, 3 KCl, 25 NaHCO3, 1 CaCl2, 7 MgCl2, 1 NaH2PO4, 20 D-glucose, 2.4 sodium pyruvate, and 1.3 sodium L-ascorbate.

The aCSF contained (in mM): 126 NaCl, 2.5 KCl, 26 NaHCO3, 1.2 CaCl2, 1.5 MgCl2, 1.3 NaH2PO4, and 10 D-glucose. Slices were incubated at 34°C before imaging.

### PI uptake assay for cell damage assessment

Acute cortical brain slices were pre-incubated in mesh-bottomed chambers containing 20 mL of carbogen oxygenated aCSF solution with 20 μM propidium iodide (PI). When blockers were used, tissue was also pre-incubated for a minimum of 20 min prior to experiment start time. Following the experiment, slices were immediately flash fixed by the SNAPSHOT method described previously^20^. Briefly, slices were fixed in PFA at 80°C for 2 minutes and then washed in 0.1 M PBS. When immunostaining was needed, slices were fixed using the same protocol, except with a glyoxal/acetic acid fixation solution^21^ : 9% (v/v) glyoxal (Sigma-Aldrich, #128465), 8% (v/v) acetic acid (Sigma-Aldrich, #695092) in PBS, pH adjusted to 4.0 using NaOH. Following fixation, slices were then permeabilized for 1-2 weeks in a tissue clearing solution comprised of 2% v/v Triton X-100 and 20% v/v DMSO dissolved in PBS (0.1 M) at 4°C prior to PI imaging. For immunostaining slices, tissue sections were first incubated in blocking solution (10% v/v donkey serum (Jackson Labs, #017-000-121) in clearing solution) for 1 day, followed by a 2 day incubation in primary antibody solution (antibody plus 2.5% v/v donkey serum in clearing solution). Primary antibody dilutions were as follows: anti-Pdgfr-β (1:200) (R&D, #AF1042), anti-Pdgfr-α (1:250) (R&D, #AF1062), anti-αSMA (1:200) (Abcam, #ab124964), anti-laminin (1:100) (Abcam, #ab11575), anti-TRPA1 (1:100) (Novus, #NB110-40763), anti-Gfap (1:200) (Invitrogen, #13-0300). Slices were washed and then incubated in secondary antibody solution (diluted to 1:200 in 2.5% v/v donkey serum in clearing solution) for 2 days. Next, slices were washed and stained with DAPI for 10 min. The secondary antibodies used were: donkey anti-goat A488 (Invitrogen, #A-11055), donkey anti-rabbit A488 (Invitrogen, #A-21206), donkey anti-goat A647 (Invitrogen, #A21447), donkey anti-rat A568 (Invitrogen, #A11057).

### Imaging of fixed and live brain slices

All imaging of either live or fixed brain tissue sections was performed using a two-photon laser-scanning microscope (LSM 7MP, Zeiss), unless stated otherwise. This microscope is equipped with a Zeiss water immersion objective lens (Plan-Apochromat 20×/numerical aperture 1.0) through which excitation and emitted light was passed. The excitation light was provided by a mode-locked Ti:Sapphire laser (Chameleon Ultra II, Coherent), which was pulsed at 80 MHz (140 fs per pulse). The excitation wavelengths for imaging were: NeuroTrace & PI, 1000 nm; A488 and PI, 800 nm. Emitted light was detected by two separate LSM BiG GaAsP detectors after passing through the appropriate emission filters. Images were acquired using line averaging (average of 8) and were processed and stored with Zen software (Zeiss). Fixed tissue was imaged free floating in PBS, and was sandwiched between a microscope slide (FisherScientific, #125442) and coverglass (Avantor, # 48393-092). Z-stack images were acquired by imaging the entire tissue volume (0-300 µm). For imaging of TRPA1 immunostaining, GAA fixed brain tissue was collected from transcardially perfused rats. 150 µm thick cortical slices were prepared, immunolabelled, and imaged on a Zeiss LSM 980 confocal microscope.

For all live tissue imaging, acute brain slices were transferred from incubation chambers to a two-photon microscope recording chamber and perfused with aCSF at 2 –3 mL/min. aCSF was bubbled with 95% O_2_/5% CO_2_ and warmed to 32 °C with a stage heater (Luigs & Neumann). For live imaging during oxidative stress induction, brain slices were first labelled by incubating brain slices in aCSF containing NeuroTrace (1:1000 dilution for 40 min). The slices were then transferred to the recording chamber and perfused with aCSF containing PI (20 µM). For testing the impacts of TRPA1 blockers on Neurotrace uptake, slices were first transferred to incubation chambers in normal aCSF or aCSF plus TRPA1 antagonists for 20 min. NeuroTrace (1:1000 dilution) was then added to the chamber and allowed to incubate for 40 min. Tissue slices were then washed and transferred to the recording chamber. Z-stack images were produced by imaging between 0 and -120 µm below the cut surface. For accurate comparison, the same gain and laser intensity were used across different brain slices.

For live pericyte Ca^2+^ imaging experiments, acute cortical brain slices from ATP13a5-CreERT2-tdTomato mice were imaged using a Zeiss LSM 510 laser-scanning confocal microscope equipped with a 40× water-immersion objective (0.8 NA). Pericytes were identified using TdTomato and GCaMP fluorescence. GCaMP6f was excited using a 488 nm laser and fluorescence was detected through a 505–530 nm bandpass filter, while TdTomato was excited using a 543 nm laser and fluorescence was detected through a 565–615 nm bandpass filter.

Pericytes were imaged approximately 15–35 µm below the cut surface of the slice. To monitor slowly developing changes in intracellular Ca^2+^ over time, image sequences containing one pericyte per slice were acquired every 3 min for up to 1 h. Brain slices were continuously perfused with aCSF at 2 mL/min and maintained at 32 ± 2 °C. In control experiments, slices were perfused with DMSO-containing aCSF for 124 minutes, followed by application of 1 mM H_2_O_2_. For TRPA1 inhibition experiments, slices were pre-treated with A-967079 (20 µM) for 14 min prior to H_2_O_2_ application, and A-967079 remained present throughout the remainder of the experiment.

### Imaging analysis

Quantification of PI positive cells was done with the following methods. Z-stack images were opened in Fiji (ImageJ) and maximum intensity projection images formed from the middle 100 µm of the 300 µm thick tissue volume. Next, a threshold fluorescence value was calculated to detect PI positive cells, and was determined as follows. For each image, the average background fluorescence was first determined by measuring the fluorescence of cells which were not labelled with PI. Then, the fluorescence threshold (F*_threshold_*) was calculated by measuring the average fluorescence of the background cell labelling (*µ_bg_*) plus five standard deviations of the sample background intensity measurements (*σ*_bg_):

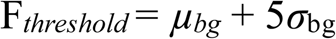

Cells which showed average somatic fluorescence at or above this threshold were categorized as PI labelled cells. For some data sets, we identified and quantified PI positive cells using ilastik (v1.4.1post1), a machine learning analysis tool for image segmentation, object classification, and cell counting. This allowed for improved speed of image analysis. Z-stack images of PI and immunolabelling fluorescence were imported into ilastik and then processed using a Pixel Classification to Object Classification workflow. Program training was performed on the Pixel Classification and Object Classification workflows, using several representative z stacks to identify ‘PI positive’ or ‘PI negative’ cells. Training was performed iteratively on several distinct Z-stacks until pixel and object classification accurately detected and classified cells across multiple images not previously used for training. In Fiji, identified PI labelled cells were categorized according to their location and/or marker expression. Cells were first categorized as vascular or non-vascular based on the spatial association between the PI labelled cell and vessels. In the presence of vascular labelling (e.g. Pdgfr-β immunolabelling or Tomato-lectin 488 (Invitrogen, #L32470)), association of PI labelled cells with vessels was readily determined. In tissue that lacked any vascular markers, vessels could be identified from transmitted light images and background PI fluorescence, which outlined vessel structures. For some experiments, PI labelled vascular cells were also categorized as being associated with primary branch order vessels (0°) or branch order vessels ≥1°. Primary order vessels were identified by their large diameter and connection to the pial surface. Vessels of branch order ≥1° were identified by their branching from a primary order vessel. In the absence of overt connection to a primary order vessel, branches were determined to be of order ≥1° based on their comparatively small diameter. When cell identification was determined with immunostaining, vascular PI labelled cells were categorized according to the presence or absence of intense labelling of Pdgfr-β, Pdgfr-α, or αSMA around the PI labelled nuclei. In Pdgfr-β expressing cells, blebs were identified as soma-associated membrane protrusions that extended at least 2 µm away from the cell body. The size of the bleb was calculated by measuring the linear distance between the farthest displacement of the bleb and the adjacent cell body or capillary wall.

For quantifying pericyte NT uptake in the presence or absence of TRPA1 blockers, Z-stack images were used to make a maximum intensity projection image in Fiji. This image represented the volume of tissue between -50 and -110 µm below the slice cut surface. Next, we calculated a threshold value, which was used to create a binary image of the NeuroTrace fluorescence. The threshold value was calculated as 20 times the average fluorescence of the background average. The binary image was then used to create a region of interest (ROI) representing all regions above the calculated threshold. The ROI was then placed on the original image, and the integrated density value (mm^2^ x F) for the NeuroTrace positive region was calculated. For live imaging of PI uptake into pericytes during oxidative stress induction, the following procedure was used for analysis. PI fluorescence was measured for the duration of the experiment from all NeuroTrace labelled pericyte cell bodies. Pericyte PI fluorescence was then background corrected by measuring time matched PI fluorescence from a neighboring region that lacked pericytes. A fluorescence threshold (F*_threshold_*) was then calculated for quantifying if a pericyte was loaded with PI. The PI loading threshold was set as 3 standard deviations of the fluorescence values measured during the baseline period (*σ_Fo_*), prior to oxidative stress induction.

For analysis of pericyte Ca^2+^, GCaMP6f fluorescence was quantified in Fiji (ImageJ). ROIs were manually traced around individual pericyte somata and maintained throughout the imaging time series. When minor displacement occurred between recordings, ROIs were manually repositioned as needed to remain accurately aligned with the same pericyte soma and corresponding background region. For each pericyte, a separate background ROI was selected in a nearby region devoid of pericyte fluorescence. Mean GCaMP6f fluorescence intensity was measured within both the somatic and background ROIs at each imaging time point.

Background-corrected pericyte fluorescence was calculated by subtracting the time-matched mean background fluorescence from the mean somatic fluorescence. Fluorescence values were then normalized to a baseline period (0–9 min), defined as the average background-corrected fluorescence measured during the first four baseline recordings. Normalized fluorescence intensity was expressed as a percentage of baseline according to:

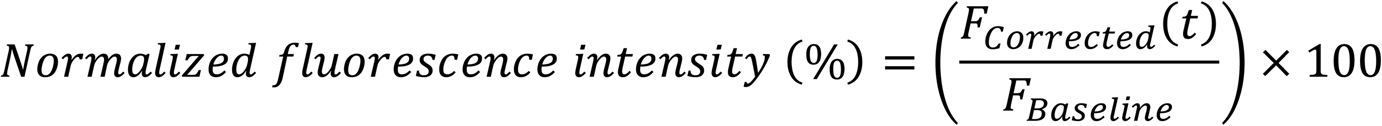

where *F_Corrected_*(*t*) represents the background-corrected somatic fluorescence at each time point and *F_Baseline_* represents the mean background-corrected fluorescence during the 0–9 min baseline period. For group analyses, only recordings up to 45 min were included to maintain a consistent number of pericyte observations across time points.

### Cell culture

Human brain vascular pericytes (ScienCell, #1200) and rat brain vascular pericytes (ScienCell, #R1200) were cultured on 75 cm^2^ flasks (ThermoFisher, #156340) using pericyte media (ScienCell, #1201) composed of: 500 ml of basal medium, 10 ml of FBS (ScienCell Cat. #0010), 5 ml of Pericyte Growth Supplement (ScienCell, Cat. #1252) and 5 ml of penicillin/streptomycin solution (ScienCell, Cat. #0503). Cells were grown until ∼90% confluent and were then detached using Accutase (StemCell Technologies, #07920). For passaging, medium was removed, and Accutase added to the flask, followed by incubation at 37°C for ∼5 min. Detached cells were collected, centrifuged at 1000 rpm for 5 min, and resuspended in fresh pericyte media. The suspension was passed through a 40 μm cell strainer and plated in new flasks. Pericytes were passaged up to a maximum of 9 times prior to use for lysate collection and western blotting.

To obtain primary rat astrocyte cultures, Sprague–Dawley rat pups (P1-P7) were used. The cortex and hippocampus were isolated, triturated with a pestle in warmed astrocyte culture medium (MEM (Gibco, #11095080) supplemented with 10% FBS (Gibco, #12483020), 0.6% glucose (Sigma-Aldrich, #G8270), 100 U/mL penicillin and 0.1 mg/mL streptomycin (Gibco, #15140122)), and further dissociated by pipetting. The tissue was incubated at 37°C for 15 min in HBSS solution (Gibco, #14175095) containing 0.25% trypsin (Gibco, #25200072). The cell suspension was filtered through a 40 μm sterile cell strainer and centrifuged to collect single cells. Cells were resuspended in astrocyte culture medium and plated in 75 cm^2^ flasks, allowing them to grow confluent for 1–2 weeks. For passaging, medium was removed, and cells were washed with PBS. 0.25% trypsin was added to cover the cells, followed by incubation at 37°C for approximately 3 min. Detached cells were collected, centrifuged at 1000 rpm for 3 min, and resuspended in fresh astrocyte culture medium. The suspension was passed through a 40 μm cell strainer and plated in new flasks or for cell lysate collection and western blotting.

### Western blotting

Primary cell cultures were harvested and homogenized on ice using RIPA lysis buffer (#89900, Thermo Scientific, Waltham, USA) supplemented with cOmplete ultra protease inhibitor tablets (#5892791001, Sigma-Aldrich) and PhosSTOP phosphatase inhibitor tablets (#4906845001, SigmaAldrich). The homogenates were centrifuged at 12,000 rpm for 10 min at 4°C to collect the supernatant. Protein concentration was then determined using the DC Protein Assay kit (#5000112, Bio-Rad, Hercules, USA). Equal protein amounts were mixed with 4× Laemmli Sample Buffer (#1610747, Bio-Rad) containing β-mercaptoethanol and vortexed at room temperature for 1h before being loaded onto Mini-Protean TGX gels (#4561096, Bio-Rad). Subsequently, SDS-PAGE was conducted, and the separated proteins were transferred onto 0.2 μm pore-size PVDF membranes using the Trans-Blot Turbo semi-dry transfer system (Bio-Rad). The membranes were then blocked for 1h at RT with 3% BSA in TBS containing 0.1% Tween-20 (TBST) and then incubated overnight at 4°C in TBST with primary antibodies. Primary antibodies used were: TRPA1 (1:300 dilution) (Novus, #NB110-40763) and GAPDH (1:1500 dilution) (Invitrogen, #PA1988). After washing, membranes were probed with HRP-conjugated secondary antibodies (Jackson ImmunoResearch, West Grove, USA) for 2h at room temperature. Protein bands were visualized using Bio-Rad Clarity Western ECL substrate and imaged with a ChemiDoc MP imaging system. Representative full-membrane western blot images are presented in Figure S2.

### Animal perfusions

To collect naïve brain tissue for immunostaining, rat brains were fixed and isolated using transcardiac perfusion with GAA. Brains were then cut into 150 µm thick coronal sections using a vibratome, as described above. Brain sections were then processed for immunostaining.

### Chemicals and reagents

Drugs and dyes used for experiments are listed in final concentration (in mM): 0.02 propidium iodide (PI) (Invitrogen, # P1304MP), 0.01 necrostatin-1 (nec-1) (Tocris, #2324/10), 0.05-0.1 z-VAD FMK (Cayman Chemical, # 14463-1), 2 diamide (Sigma Aldrich, #D3648), 1 H_2_O_2_ (Sigma Aldrich, #H1009), 0.03 cyclopiazonic acid (CPA) (Sigma Aldrich, #C1530), 0.1 NiCl_2_ (Sigma), 0.05 CdCl_2_, 0.1 SKF-96365, 0.02 SET2, 0.02 ruthenium red (RR), 1-2 amiloride, 0.2-0.4 benzamil, 0.02 A-967079 (A96), 0.01 HC-030031 (HC-03) (Sigma Aldrich, 557440). All drugs or chemicals were dissolved into the aCSF. DMSO was used as the vehicle to dissolve A96, HC-03, necrostatin-1, z-VAD FMK, CPA and SET2. DMSO was diluted to a maximal working concentration of 0.1% (v/v).

### Statistics and reproducibility

All summary graph data shown represent the mean ± standard error of mean. For violin plots, medians and quartiles are depicted within each graph. For each data set, the slice replicates and the animal replicates are described within the summary data and figure legends. The slice replicate number is represented by the ‘n’ value and the animal replicate number by the ‘N’ value. The Shapiro-Wilk normality test was used to examine data sets for normality. If data were normal, a Student’s paired or unpaired t test was used to test for differences between two means, while a one-way ANOVA with a Dunnett’s multiple comparisons test was used to test for differences between multiple means and a control condition. If the data were not normally distributed, a Mann–Whitney U test was used for two means, whereas a Kruskal-Wallis test with Dunn’s multiple comparison test was used to compare multiple means to a control condition. A Welch’s t test was used to compare two data sets if the variances differed significantly. For paired data sets, a paired t test was used when the data was normally distributed, while a Wilcoxon matched-pairs signed rank test was used when the data sets were not normally distributed. Tests were considered statistically significantly different at *p* ≤ 0.05. The exact p-values are indicated in the figure legends. All data sets presented were produced from distinct experiments, with the exception of Figure 4b,c. The data used for this analysis was also used for a distinct analysis in Figure 2b,c.

## Results

We first examined the impact of acute oxidative stress on tissue damage in rodent brain slices. Acute rat cortical slices were incubated in artificial cerebrospinal fluid (aCSF) containing pro-oxidants and propidium iodide (PI, 20 µM) (Figure 1a). Cellular PI uptake was used to assess cell damage, as it reflects membrane breakdown or the opening of large-pore channels engaged during cell dysfunction ^14,22–24^. We first treated tissue with hydrogen peroxide (H_2_O_2_), an endogenous ROS generated through the dismutation of superoxide ^5^. Under physiological conditions, extracellular H_2_O_2_ is estimated to be in the low micromolar range, but it can become significantly elevated during pathological conditions such as ischemia-reperfusion ^25^. Following incubation with a high concentration of H_2_O_2_ (1 mM) for 1 hour, brain slices were flash fixed ^20^ and PI labelling was imaged with two-photon microscopy, as described previoussly ^14^. Exposure to H_2_O_2_ produced a significant increase in the number of PI-labelled cells relative to control conditions (Figure 1b). Because oxidation of protein thiols is a major downstream action of ROS ^26–28^, we also examined cell damage following tissue exposure to diamide, which oxidizes the cellular thiol pool located on proteins and glutathione ^29^. We have previously shown that diamide is a robust inducer of cellular oxidation in brain tissue ^26^. Exposure of brain slices to diamide (2 mM) for 1 hour resulted in a significant increase in PI-labelled cells in the cortical tissue (Figure 1b). Consistent with the known neuronal sensitivity to oxidative stress^13,18^, we observed PI-labelled neurons in the upper cortical layers (Figure 1b). However, we also observed an abundance of small, non-neuronal PI-labelled cells in tissue exposed to either H_2_O_2_ or diamide (Figure 1b). Visualization of the microvasculature revealed that this cell population was closely associated with the small-diameter capillaries as well as primary branch-order vessels connecting to the pial surface (Figure 1c). Quantification confirmed a significant increase in PI-labelled microvascular cells following exposure to either H_2_O_2_ or diamide (Figure 1d). In the non-vascular parenchyma, oxidative stress evoked by H_2_O_2_, but not diamide, significantly increased the number of damaged, PI-labelled cells (Figure 1d). This discrepancy may be attributed to mechanistic differences between these pro-oxidants; unlike diamide, H_2_O_2_ can directly drive lipid and nucleic acid oxidation via hydroxyl radical generation ^5,29^. When comparing the relative proportions of damaged vascular versus non-vascular cells, in H_2_O_2_ there was no significant difference, whereas after diamide exposure there was a significantly greater percentage of damaged cells associated with the vasculature (Figure 1d). Considering that vascular cells comprise only a minority of the total cell population in cortical layers I/II ^30^, this high proportion of damaged vascular cells suggests a comparatively high sensitivity to oxidative stress in the microvasculature.

**Figure 1.**
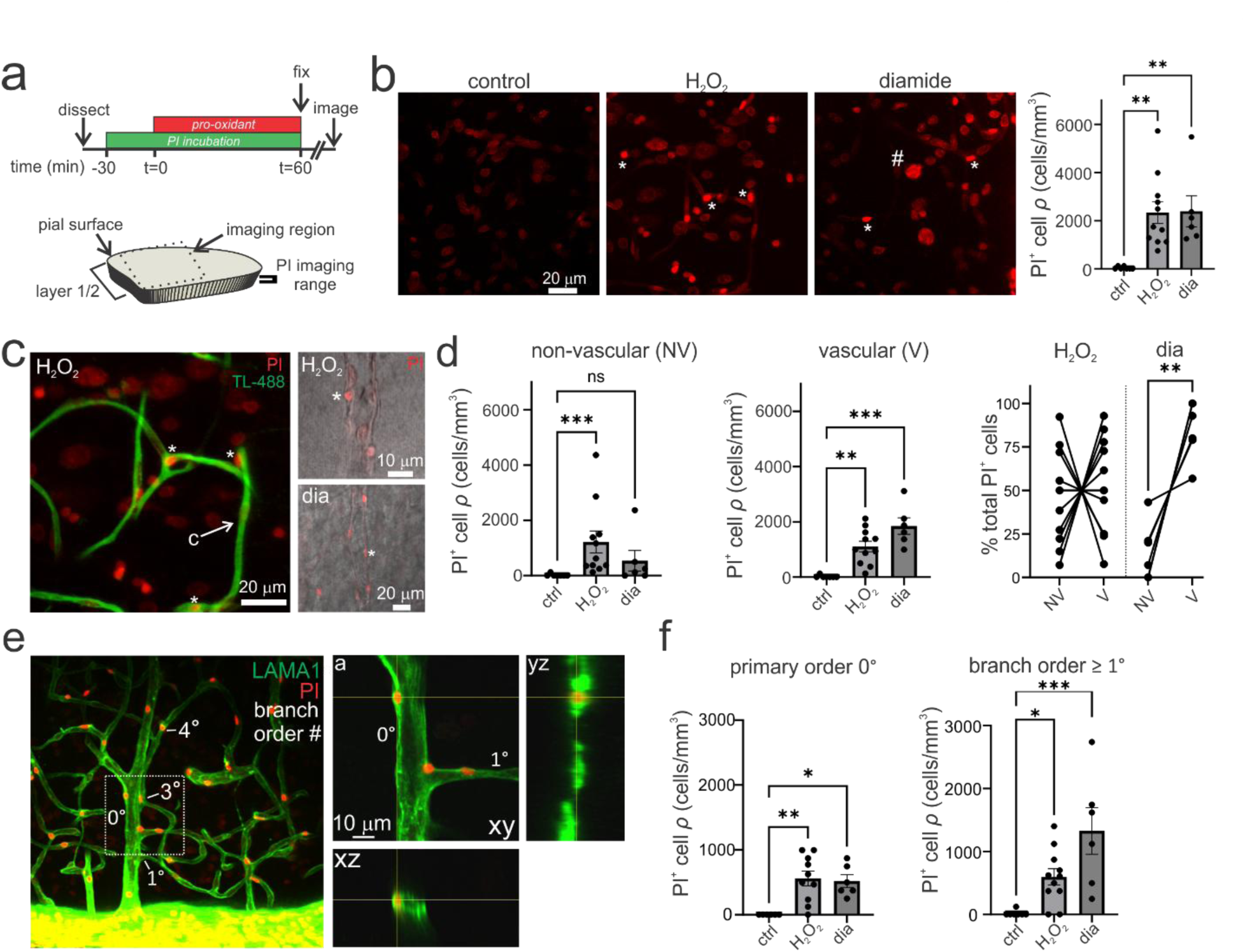
Oxidative stress causes damage to vascular cells in multiple microvascular zones. **a)** Illustration showing the timeline of experiments during the induction of oxidative stress and measurement of propidium iodide (PI) uptake. The region imaged for analysis is the upper cortical layers (1/2) and the healthy mid-section of the acute brain slice. **b)** *Left*, Induction of oxidative stress with H_2_O_2_ (1 mM) or diamide (dia) (2 mM) cause PI uptake in a fraction of cells in the cortical tissue. Images reflect maximum intensity projections Asterisks (*) denote small, PI labelled (PI^+^) cells and hash (*#*) denote a large, PI labelled neuron. H_2_O_2_ or diamide cause a significant increase in the abundance of PI labelled cells per volume of tissue, relative to control (control: *n*=7, *N*=5; H_2_O_2_: *n*=11, *N*=6; dia: *n*=6, *N*=4; Kruskal Wallis test: df=2, *H*=14.44, P=0.0007; Dunn’s multiple comparisons test: control vs H_2_O_2_ (P=0.001), control vs dia (P=0.003). **c)** Oxidative stress causes the appearance of PI labelled cells on tomato lectin (TL-488) stained capillaries (c) (*Left*) and large caliber primary order vessels, visualized with transmitted light imaging (*Right*). **d)** *Left*, Oxidative stress induction with H_2_O_2_, but not diamide, causes a significant increase in the density of non-vascular PI labelled cells relative to control (control: *n*=7, *N*=5; H_2_O_2_: *n*=11, *N*=6; dia: *n*=6, *N*=4; Kruskal Wallis test: df=2, *H*=13.49, P=0.0012; Dunn’s multiple comparisons test: control vs H_2_O_2_ (p=0.0005), control vs dia (p=0.17)). *Middle*, H_2_O_2_ or diamide cause a significant increase in the abundance of PI labelled, vessel-associated cells relative to control (control: *n*=7, *N*=5; H_2_O_2_: *n*=11, *N*=6; dia: *n*=6, *N*=4; Kruskal Wallis test: df=3, *H*=16.03, P=0.0003; Dunn’s multiple comparisons test: control vs H_2_O_2_ (p=0.0041), control vs dia (P=0.0003). *Right*, In H_2_O_2_, the percentage of total PI labelled cells that are vascular (V) is not significantly greater than the proportion of non-vascular (NV) cells (non-vascular: *n*=11, *N*=6; vascular: *n*=11, *N*=6; paired t-test: df=10, t=0.472, P=0.65). In diamide, there is a significantly greater percentage of PI labelled cells associated with the vasculature vs non-vascular regions (non-vascular: *n*=6, *N*=4; vascular: *n*=6, *N*=4; paired t-test: df=5, t=5.15, P=0.0036). **e)** *Left*, Maximum intensity projection image of the cortical microvasculature immunolabelled for laminin shows that PI labelled cells are found on both primary order vessels (0°) and vessels of branch orders ≥1°. *Right*, PI labelled cells are embedded in the laminin labelled basement membrane. **f)** Oxidative stress induced by H_2_O_2_ or diamide cause a significant increase in PI labelled cell abundance on primary order (0°) and branch order vessels ≥1°. Primary order vessels (control: *n*=6, *N*=4; H_2_O_2_: *n*=10, *N*=5; dia: *n*=6, *N*=4; Kruskal Wallis test: df=2, *H*=11.41, P=0.001; Dunn’s multiple comparisons test: control vs H_2_O_2_ (P=0.0028), control vs dia (P=0.0143)). Branch orders ≥1° (control: *n*=7, *N*=5; H_2_O_2_: *n*=11, *N*=6; dia: *n*=6, *N*=4; Kruskal Wallis test: df=2, *H*=13.01, P=0.0015; Dunn’s multiple comparisons test: control vs H_2_O_2_ (P=0.0194), control vs dia (P=0.0009)). Graphs depict the mean ± SEM. Each data point represents a single brain slice replicate. The number of brain slice replicates is represented by the ‘n’ value. The number of animal replicates is represented by the ‘N’ value. P values: *P≤0.05, **P≤0.01, ***P≤0.001, ****P≤0.0001.

The cortical microvasculature is comprised of discrete zones, including penetrating arterioles, arteriole-capillary transitions (ACT), the mid-capillary bed, post-capillary venules, and ascending venules ^31^. PI-labelled cells were clearly observable on primary-order vessels and on early branches from these vessel segments (Figure 1e). Numerous PI-labelled cells were also detected on mid-capillary bed vessels, but branch orders could not always be definitively determined. As such, we categorized cells according to their location on primary-order vessels (0°: penetrating arterioles and ascending venules) or branches of primary-order vessels (branch order ≥ 1°: ACT, mid-capillary bed, and post-capillary venules). Quantification revealed that oxidative stress evoked by either H_2_O_2_ or diamide produced a significant increase in PI-labelled cells on both primary-order branches and branches of order ≥ 1° (Figure 1f). Furthermore, laminin staining confirmed that damaged vessel-associated cells were located within the vascular basement membrane, consistent with a vascular cell population rather than vessel-associated glia (Figure 1e). These data demonstrate the presence of an oxidative stress-sensitive population of vascular cells across multiple microvascular zones.

Next, we explored the identity of this sensitive vascular cell population. The microvasculature cell population includes endothelial cells, smooth muscle cells, pericytes, and perivascular fibroblasts ^31,32^. On branches of order ≥ 1°, representing the ACT, mid-capillary bed, and post-capillary venules, the vascular wall consists primarily of endothelial cells and pericytes ^32^. Due to their “bump-on-a-log” morphology, we hypothesized that the damaged cells on these branches were pericytes, which can be distinguished from endothelial cells by the expression of Pdgfr-β ^31^. Because standard PFA fixation yielded poor immunolabelling across thick tissue slices (Figure S1a), we used an alternative fixation method with glyoxal acetic acid (GAA), which has been reported to improve the labelling of physically inaccessible synaptic proteins in brain tissue ^21^. GAA fixation significantly improved the depth-resolved immunolabelling of Pdgfr-β and alpha-SMA relative to PFA (Figure S1a,b), allowing reliable identification of Pdgfr-β-positive cells throughout the tissue volume. On vessel branches of order ≥ 1°, PI-labelled cells in H_2_O_2_ - and diamide-treated slices expressed Pdgfr-β (Figure 2a), identifying them as pericytes. Notably, under oxidative stress, many PI-labelled pericytes displayed pronounced membrane blebbing protruding from the cell body into the parenchymal space (Figure 2a), a morphology consistent with pericyte stress and damage ^31,33^. Interestingly, adjacent Pdgfr-β negative endothelial cells rarely exhibited PI uptake (Figure 2a). In addition to vessel branches of order ≥ 1°, many PI-labelled cells on primary-order vessels in diamide-treated tissue also expressed Pdgfr-β (Figure 2a). Analysis across all microvascular zones confirmed that oxidative stress induced by either H_2_O_2_ or diamide significantly increased the abundance of PI-labelled Pdgfr-β expressing cells (Figure 2b). Vessel branches of order ≥ 1° showed a significant increase in PI-positive Pdgfr-β pericytes after treatment with either pro-oxidant (Figure 2b). On primary-order vessels, diamide, but not H_2_O_2_, significantly increased the abundance of PI-labelled Pdgfr-β cells (Figure 2b), likely reflecting differences in cellular antioxidant buffering against hydrogen peroxide versus direct thiol modifiers ^29^.

**Figure 2.**
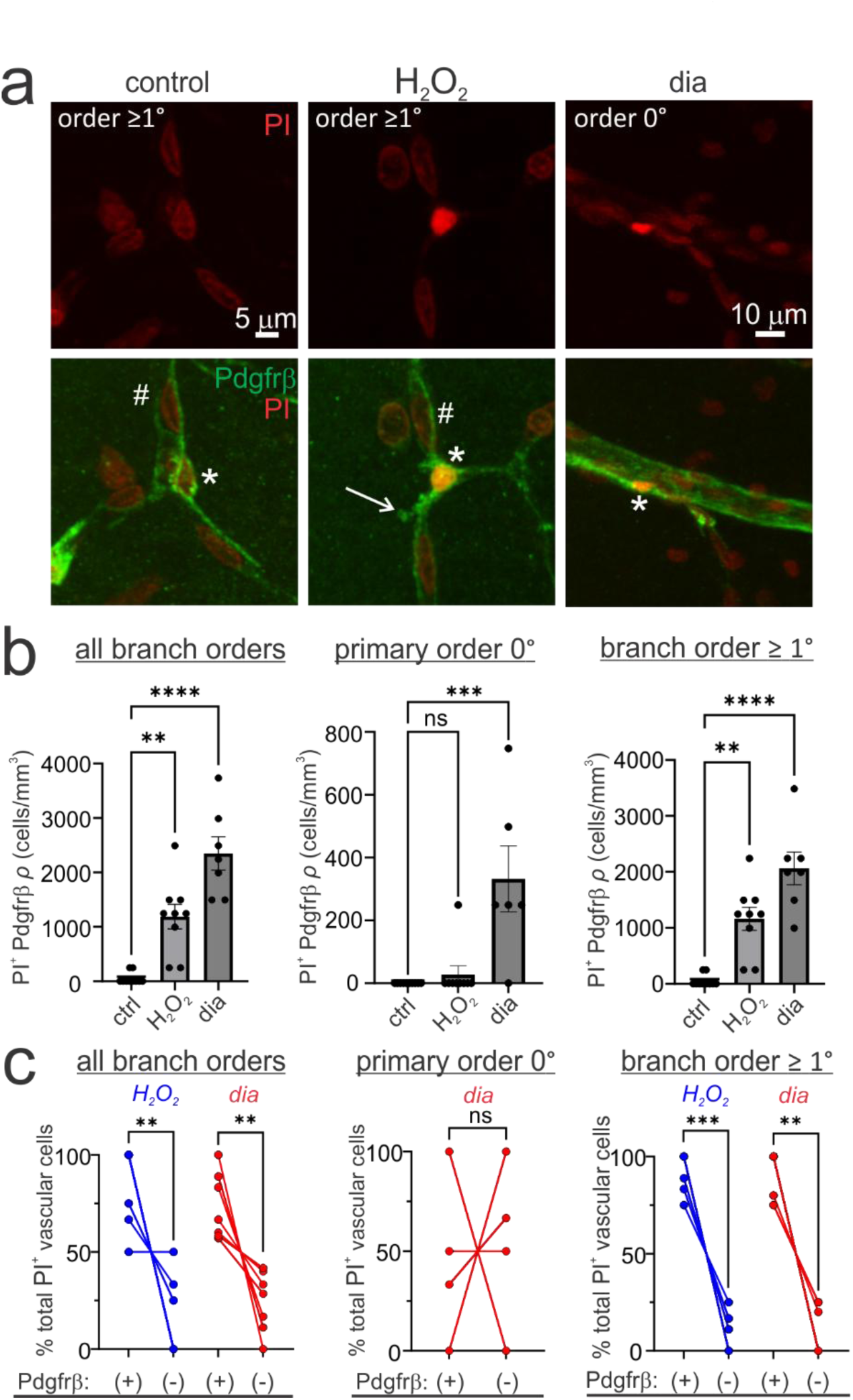
Pdgfr-β expressing microvascular cells are sensitive to oxidative stress. **a)** *Left*, Image depicts a Pdgfr-β immunolabelled pericyte on a capillary (branch order ≥1°) in control conditions. *Middle*, A PI labelled pericyte in H_2_O_2_ (1 mM) showing a prominent bleb structure near the cell body (*arrow*). *Right*, A PI labelled, Pdgfr-β expressing cell on a primary order vessel (branch order 0°) after oxidative stress induction with diamide (2 mM). Asterisk (*) denotes Pdgfr-β positive cells and hash (#) denotes Pdgfr-β negative endothelial cells. **b)** *Left*, Summary data showing that H_2_O_2_ or diamide significantly increase the number of PI labelled Pdgfr-β positive cells measured across all microvascular branch orders (control: *n*=10, *N*=3; H_2_O_2_: *n*=9, *N*=3; dia: *n*=7, *N*=2; Kruskal Wallis test: df=2, *H*=20.57, P<0.0001; Dunn’s multiple comparisons test: control vs H_2_O_2_ (P=0.009), control vs diamide (P<0.0001). *Middle*, Oxidative stress induced with diamide, but not H_2_O_2_, increases the number of PI labelled Pdgfr-β expressing cells on primary order vessels (control: *n*=10, *N*=3; H_2_O_2_: *n*=9, *N*=3; dia: *n*=6, *N*=2; Kruskal Wallis test: df=2, *H*=15.29, P=0.0005; Dunn’s multiple comparisons test: control vs H_2_O_2_ (P>0.9999), control vs diamide (P=0.0004). *Right*, H_2_O_2_ or diamide produce a significant increase in PI labelled Pdgfr-β expressing cells on vessel branch orders ≥1° (control: *n*=10, *N*=3; H_2_O_2_: *n*=9, *N*=3; dia: *n*=7, *N*=2; Kruskal Wallis test: df=2, *H*=19.51, P<0.0001; Dunn’s multiple comparisons test: control vs H_2_O_2_ (P=0.0054), control vs diamide (P<0.0001. **c)** *Left*, In H_2_O_2_ or diamide treated tissue, the proportion of PI labelled vascular cells from all microvascular branch orders which express Pdgfr-β (+) is significantly greater than the proportion of PI labelled cells not expressing Pdgfr-β (-) (H_2_O_2_: *n*=9, *N*=3; Wilcoxon matched pairs signed rank test, *W*=-36, P=0.0078; diamide: *n*=7, *N*=2; Wilcoxon matched pairs signed rank test, *W*=-28, P=0.0156). *Middle*, In diamide treated tissue, there was no significant difference in the proportion of PI labelled cells which expressed or did not express Pdgfr-β on primary order vessels (paired t test, t=0.916, df=5, P=0.4). *Right*, For branch orders ≥1°, in H_2_O_2_ or diamide, there was a significantly greater percentage of PI labelled cells which expressed Pdgfr-β than the proportion that did not (H_2_O_2_: *n*=9, *N*=3; Wilcoxon matched pairs signed rank test, *W*=45, P=0.004; diamide: *n*=7, *N*=2; Wilcoxon matched pairs signed rank test, *W*=28, P=0.0156).

We next quantified the relative proportions of PI-labelled cells that did or did not express Pdgfr-β within each vascular zone. Across all microvascular branches, there was a significantly higher percentage of PI-labelled cells that expressed Pdgfr-β compared to those that did not (Figure 2c). On primary-order vessels from diamide-treated tissue, the proportion of PI-labelled Pdgfr-β cells was not significantly different from Pdgfr-β cells (Figure 2c). On branches of order ≥ 1°, however, Pdgfr-β -expressing cells accounted for the vast majority of all PI-labelled cells (Figure 2c). These findings indicate that the majority of damaged microvascular cells under oxidative stress express Pdgfr-β, and that capillary pericytes are particularly sensitive relative to adjacent endothelial cells.

To further validate the sensitivity of pericytes to oxidative stress, we imaged pericytes and PI uptake during pro-oxidant treatment using live two-photon imaging in acute brain slices. To do this, pericytes were stained with NeuroTrace 500/525, a reliable and selective live cell pericyte marker ^34^. Tissue slices were then perfused with aCSF containing extracellular PI, to label cell damage, and oxidative stress was triggered with diamide application. Induction of oxidative stress with diamide resulted in the appearance of PI fluorescence in a high percentage of NeuroTrace labelled pericytes within 20 minutes (Figure 3a,b). Interestingly, there was also a simultaneous loss of NeuroTrace fluorescence from cells presenting with PI uptake (Figure 3a). The simultaneous uptake of PI and loss of NeuroTrace is consistent with a rapid and progressive cell membrane disruption. As such, these results provide further evidence that pericytes are highly sensitive to oxidative stress.

**Figure 3.**
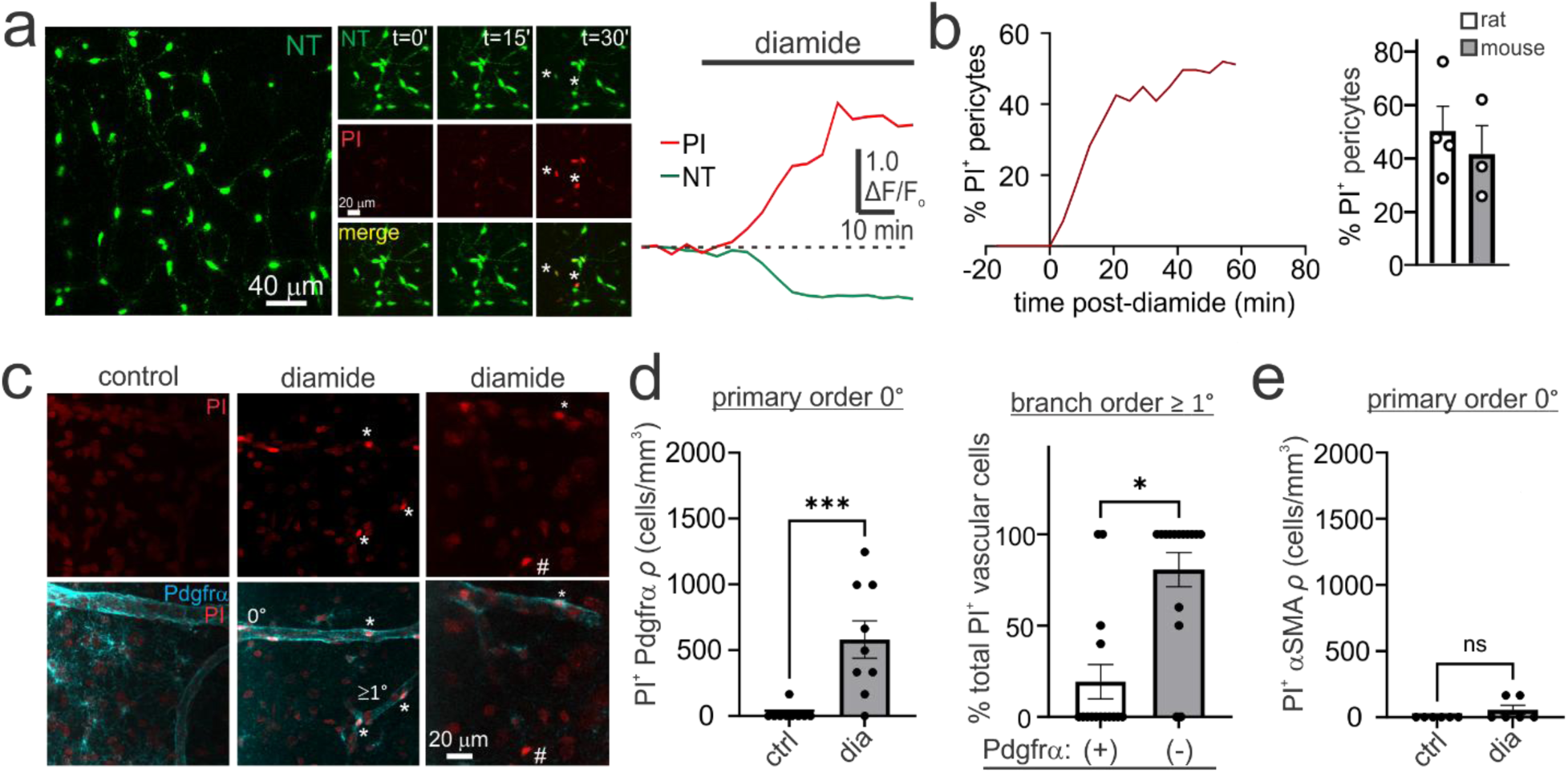
Pericytes and perivascular fibroblasts are oxidative-stress sensitive microvascular cells. **a)** *Left*, Live imaging of pericytes in an acute cortical brain slice using NeuroTrace (NT) labelling. Simultaneous measurement of NT and PI fluorescence during the induction of oxidative stress with diamide (2 mM). Following several minutes of diamide exposure, PI fluorescence intensity increases and NT fluorescence decreases in pericytes. *Right*, Measurement of NT and PI fluorescence from a single pericyte reveals a simultaneous increase in PI uptake and a loss of NT fluorescence following ox-stress induction. **b)** *Left*, Plot from a single brain slice experiment showing that the percentage of PI labelled (PI^+^) pericytes in the imaging field substantially increases within minutes of ox-stress induction. *Right*, Summary data showing the mean percentage of pericytes in the slice imaging area which loaded with PI after oxidative stress induction with diamide. Experiments were performed in rat or mouse acute brain slices. **c)** Oxidative stress induction with diamide results in PI labeling of cells on vessels of primary order (0°) and branch orders ≥1° which stain for Pdgfr-α (*: Pdgfr-α positive vascular cell). Many PI labelled cells on branch orders ≥1° did not express Pdgfr-α (#: Pdgfr-α negative vascular cell). **d)** *Left*, Relative to control, oxidative stress induction with diamide significantly increases the density of PI labelled Pdgfr-α expressing cells on primary order vessels (control: n=9, *N*=3; diamide n=9, *N*=3; Mann-Whitney U test, *U=*5.5, P= 0.0008). *Right*, On branch orders ≥1°, a significantly greater percentage of PI positive cells do not express Pdgfr-α(-) compared to the proportion that express Pdgfr-α(+) (*n=*15, *N*=3; Wilcoxon matched pairs signed rank test, *W*=73, P=0.013). **e)** Oxidative stress induction with diamide did not significantly increase PI labelling in α-SMA expressing cells of primary order vessels (control: *n*=6, *N*=3; diamide: *n*=6, *N*=3; *U*=12, P=0.45).

As pericytes are absent from primary-order vessels, we next resolved which Pdgfr-β expressing cell type was damaged in this zone. Pdgfr-β expressing cells on penetrating arterioles and ascending venules are either smooth muscle cells or perivascular fibroblasts ^31,32,35^. In fixed brain slices previously treated with diamide, many PI labelled cells on primary-order vessels exhibited Pdgfr-α immunoreactivity (Figure 3c), with a significant increase in PI-positive Pdgfr-α expressing cells relative to control conditions (Figure 3d). While perivascular fibroblasts are largely absent from capillary beds, they occupy low order branches from penetrating arterioles ^32^. Indeed, we observed occasional PI labelled Pdgfr-α expressing cells on low-order branches (Figure 3c); however, Pdgfr-α expressing cells comprised only a minor fraction of the total damaged cells on branches of order ≥ 1° (Figure 3d), confirming that pericytes constitute the dominant damaged population on vessel orders ≥ 1°. Finally, diamide exposure produced no significant increase in PI labelled α-SMA expressing smooth muscle cells on primary-order vessels (Figure 3e). Thus, perivascular fibroblasts-rather than smooth muscle cells-represent the oxidative stress-sensitive Pdgfr-β expressing population on primary order vessels.

Pericytes play well-established roles in regulating capillary diameter, local blood flow, and blood-brain barrier integrity ^36–40^. Although pericyte stress and degeneration are prominent features of multiple neurological conditions, the underlying mechanisms remain poorly understood ^39,41,42^. Based on these considerations, we focused our subsequent mechanistic investigation on resolving the pathways driving pericyte vulnerability and damage. Elevated intracellular Ca^2+^ is a primary trigger of cytotoxicity ^43^. We therefore tested whether Ca^2+^ signaling mediates microvascular damage during acute oxidative stress. For initial screening efficiency across multiple pharmacological conditions, we quantified total PI labelled cells across all microvascular vessel orders collectively, as pericytes represent the dominant damaged population across the microvascular network. Remarkably, incubation in Ca^2+^-free aCSF virtually abolished PI uptake across the microvasculature in tissue exposed to either H_2_O_2_ or diamide (Figure 4a). In contrast, inhibiting canonical programmed cell death cascades with the pan-caspase inhibitor z-VAD-FMK (apoptosis) ^44^, or necrostatin-1 (necroptosis) ^45^, failed to confer protection (Figure 4a). Necrostatin-1 actually exacerbated H_2_O_2_ -induced damage, potentially due compensatory cell death signaling ^46^. These results indicate that oxidative microvascular injury does not proceed through classical programmed cell death pathways, but instead requires plasma membrane Ca^2+^ influx.

**Figure 4.**
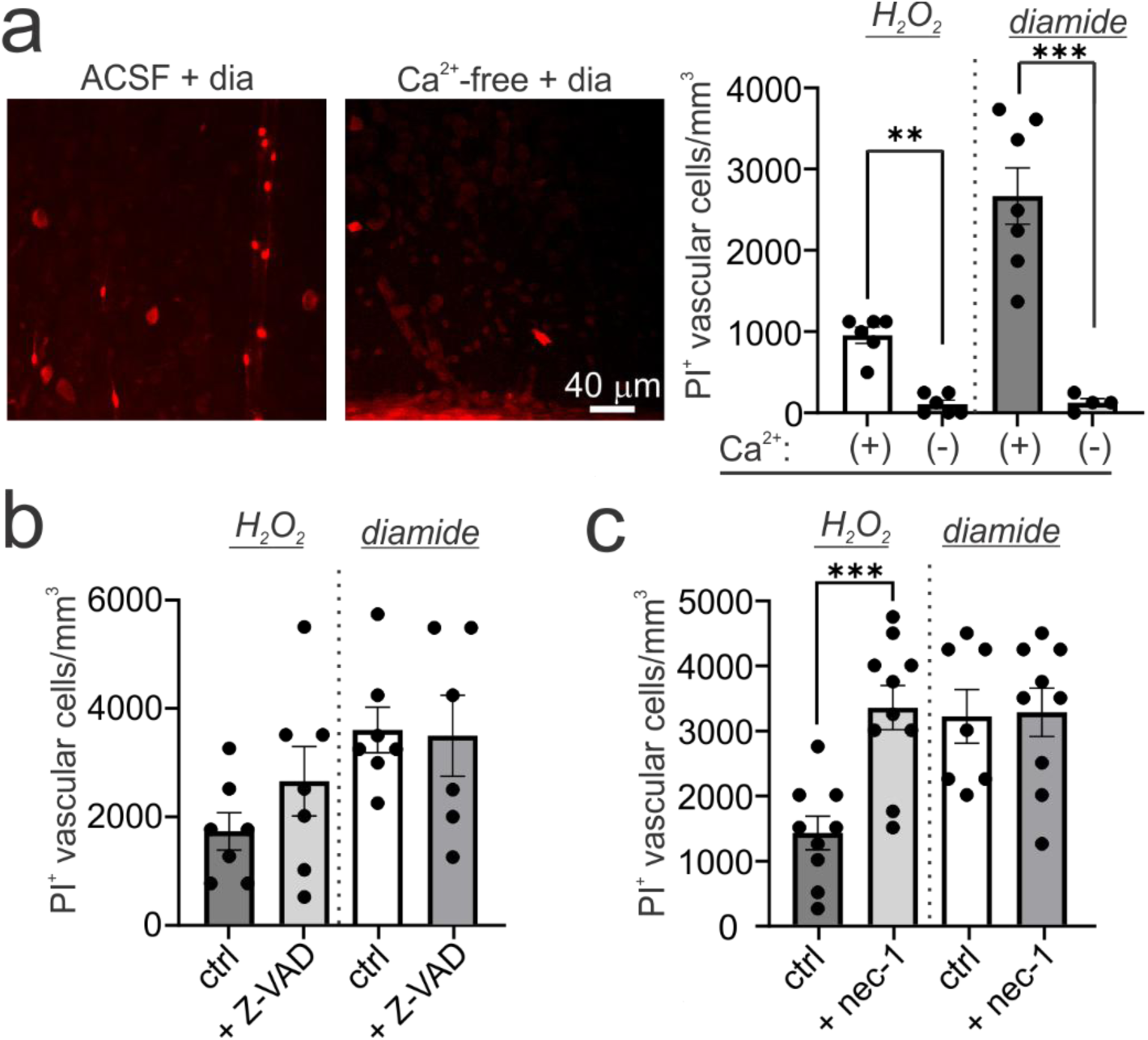
Oxidative damage of microvascular cells requires Ca^2+^ influx but not apoptotic or necroptotic mechanisms. **a)** *Left*, Images showing that the number of PI labelled vessel associated cells is substantially reduced when extracellular Ca^2+^ is excluded (Ca^2+^-free) from the ACSF during oxidative stress induction with diamide. *Right*, The density of PI labelled vascular cells in the presence of H_2_O_2_ (1 mM) or diamide (2 mM) are significantly reduced when brain tissue is incubated in Ca^2+^-free ACSF (H_2_O_2_ + ACSF: *n*=6, *N*= 3; H_2_O_2_ + Ca^2+^-free ACSF: *n*=6, *N*=2; Mann-Whitney U test, *U*=0, P=0.0022; diamide + ACSF: *n*=7, *N*=5; diamide + Ca^2+^-free ACSF *n*=4, *N*=3; unpaired student’s t test: t=5.4, df=9, P=0.0004). **b)** Pre-treating brain slices with the pan-caspase inhibitor, Z-VAD (50 & 100 µM, pooled data) had no significant impact on the number of PI labelled cells associated with the microvasculature in H_2_O_2_ or diamide (H_2_O_2_: *n*=7, *N*=2; H_2_O_2_ + Z-VAD: *n*=7, *N*=2; Welch’s t test: t=1.271, df=9.22, P=0.235; diamide: *n*=7, *N*=2; diamide + Z-VAD: *n*=6, *N*=2; Welch’s t test: t=0.1245, df=8, P=0.904). **c)** Pre-treating brain slices with the RIPK1 inhibitor, nec-1 (10 µM), in the presence of H_2_O_2_, significantly increased the density of PI labelled cells on the microvasculature. (H_2_O_2_: *n*=9, *N*=3; H_2_O_2_ + nec-1: n=10, N=3; Welch’s t test: t=4.5, df=16.29, P=0.0003). Nec-1 did not significantly impact the number of PI labelled cells induced by diamide (diamide: *n*=7, *N*=2; diamide + Z-VAD: *n*=9, *N*=3; Welch’s t test: t=0.114, df=13.2, P=0.91).

We next tested candidate Ca^2+^ entry and release pathways for their protective effects on the microvascular cells. Depleting intracellular Ca^2+^ stores with cyclopiazonic acid (CPA) ^47^ had no significant impact on diamide-evoked damage (Figure 5b). Similarly, broad blockade of voltage-gated Ca^2+^ channels with Cd^2+^ and Ni^2+^ had no protective effect on microvascular cell PI uptake (Figure 5b) ^48^. In contrast, the pan-TRP channel blocker ruthenium red (20 µM) ^49^ significantly attenuated microvascular cell damage (Figure 5a,b). Among ROS-sensitive TRP channel families ^50^, we first examined TRPC and TRPV subclasses: neither SKF-96365 (100 µM; targeting TRPC and ORAI channels) ^51,52^ nor SET2 (20 µM; targeting TRPV2) ^53^ altered microvascular cell damage (Figure 5b). Both TRP ankyrin 1 (TRPA1) and TRP polycystin channels (TRPP) are documented to be ROS-sensitive cation channels ^54,55^ . We therefore tested amiloride, which acts as a broad antagonist of both TRPA1 and TRPP channels at low millimolar concentrations ^56,57^. Pre-treatment with amiloride (2 mM) produced a potent reduction in microvascular cell damage during oxidative stress (Figure 5a,b). We next evaluated benzamil, an amiloride derivative with lower IC_50_ values than amiloride for both TRPA1 and TRPP channels ^56,57^. Pre-treatment with benzamil (200–400 µM) similarly conferred robust protection to microvascular cells (Figure 5a,b). Thus, our results implicate a Ca^2+^ entry pathway in driving microvascular cell damage that is sensitive to the broad TRP channel blocker ruthenium red, amiloride, and an amiloride-derivative.

**Figure 5.**
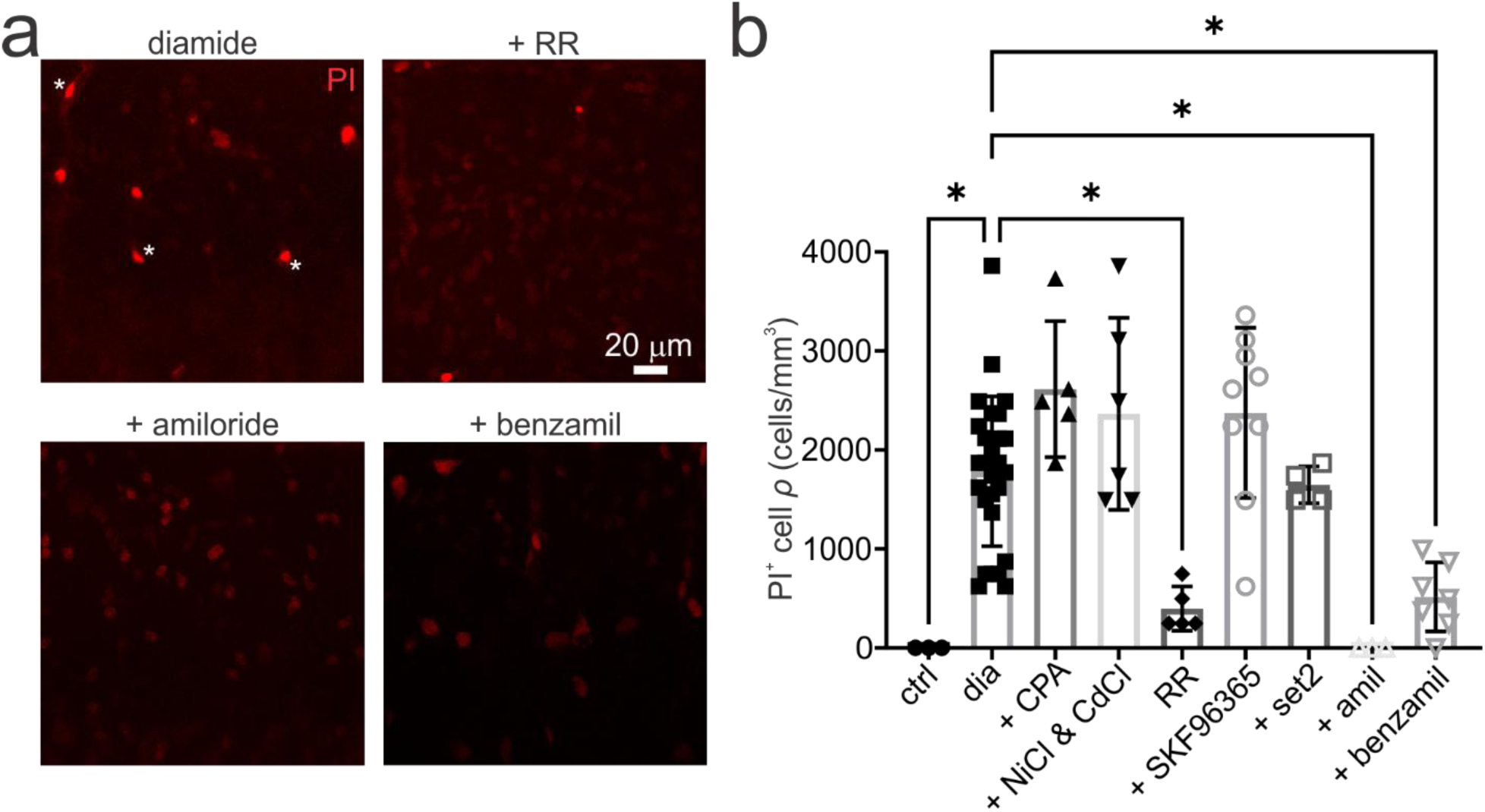
Discrete Ca^2+^ channel antagonists prevent microvascular damage during oxidative stress. **a)** Pre-treating brain slices with ruthenium red (20 µM), amiloride (2 mM), or benzamil (200 µM) substantially reduce diamide-evoked PI uptake into microvascular cells. Asterisks identify PI labelled microvascular cells. **b)** Summary data showing that the following treatments have no significant effect on diamide-evoked PI uptake into microvascular cells: depleting endoplasmic reticulum Ca^2+^ stores with CPA (30 µM), blocking voltage gated Ca^2+^ channels with Ni^2+^ (100 µM) + Cd^2+^ (50 µM), inhibiting TRPC channels with SKF96365 (100 µM), blocking TRPV2 with SET2 (20 µM). By contrast, the following treatments significantly reduce diamide-evoked PI uptake into microvascular cells: ruthenium red (20 µM), amiloride (2 mM) or benzamil (200 µM and 400 µM conditions pooled) (Kruskal-Wallis test, KW=42.27, P<0.0001; Dunn’s multiple comparisons test: diamide vs control, P= 0.0299; diamide vs CPA, P=0.82; diamide vs Ni^2+^ & Cd^2+^, P>0.999; diamide vs ruthenium red, P=0.0487; diamide vs SKF96365, P= 0.999; diamide vs set2, P=0.999; diamide vs amiloride, P=0.0299, diamide vs benzamil, P=0.0306).

TRPA1 channels are well-known to be heavily involved in tissue inflammation and ROS signaling, and have been reported to have functions in endothelial cells and astrocytes of the neurovascular unit ^58–61^. Moreover, several established TRPA1 selective antagonists exist.

Conversely, TRPP channels are not well described in the brain and selective inhibitors have not been developed. As such, we next examined the impact of selective blockade of TRPA1 on microvascular cell damage. We tested the selective TRPA1 antagonists A96 and HC-03 ^62^. Pre-treatment with A96 (20 µM) robustly protected microvascular cells during oxidative stress (Figure 6a,b). Next, we quantified the impacts of these drugs specifically on pericytes, using Pdgfr-β immunolabelling to validate cell identity in H_2_O_2_ or diamide treated tissue. Analysis revealed that the abundance of PI labelled Pdgfr-β expressing cells on the vessel branches of order ≥ 1°, which largely correspond to pericytes, was significantly reduced by pre-treatment with A96 (20 µM) or HC-03 (10 µM) (Figure 6c,d). Furthermore, A96 pre-treatment significantly reduced the number of pericytes exhibiting membrane blebbing, restoring morphology to levels similar to control conditions (Figure 6c,e). These data indicate that inhibition of TRPA1 is able to prevent pericyte structural damage during acute oxidative stress.

**Figure 6.**
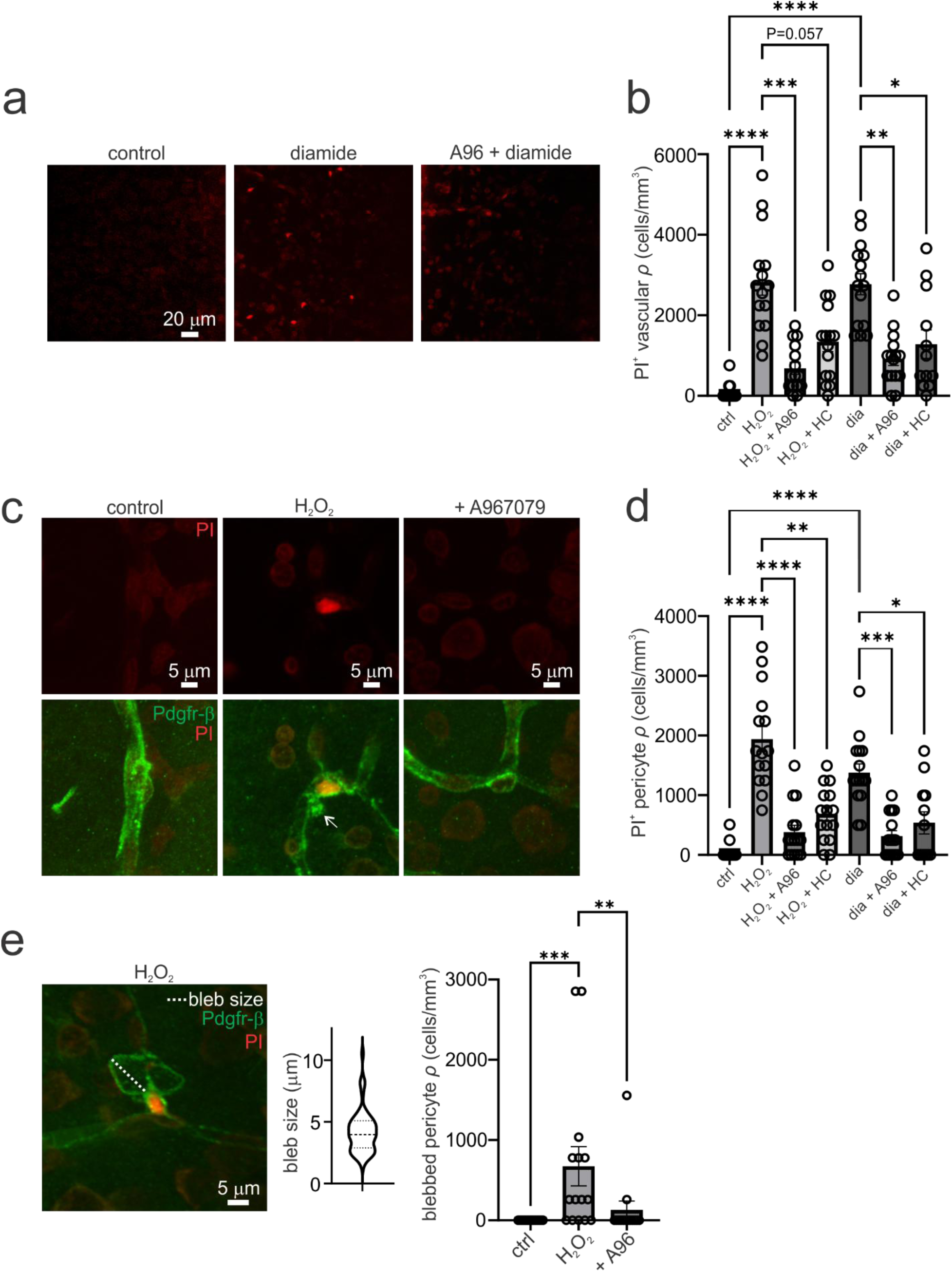
TRPA1 activity underlies pericyte damage during oxidative stress. **a)** Images of PI labelling show that pre-treatment with the TRPA1 antagonist, A96 (20 µM), substantially reduces the abundance of PI labelled microvascular cells evoked by oxidative stress in diamide (2 mM). **b)** Summary data showing that the TRPA1 antagonist, A96, significantly reduces the abundance of PI labelled microvascular cells evoked by H_2_O_2_ or diamide. Another TRPA1 antagonist, HC-03 (10 µM), significantly reduces the number of PI labelled microvascular cells in diamide. In H_2_O_2_, HC-03 causes a near significant reduction in PI labelled cells (control: *n*=15, *N*=5; H_2_O_2_: *n*=15, *N*=5; H_2_O_2_ + A96: *n=*15, *N*=5; diamide: *n*=15, *N*=5; diamide + A96: *n*=15, *N*=5; diamide + HC-03: *n=*12, *N*=4; Kruskal-Wallis test, KW=61.23, P<0.0001; Dunn’s multiple comparisons test, control vs H_2_O_2_: P<0.0001; H_2_O_2_ vs H_2_O_2_ + A96: P= 0.0002; H_2_O_2_ vs H_2_O_2_ + HC-03: P=0.0577; control vs diamide: P<0.0001; diamide vs diamide + A96: P=0.0019; diamide vs diamide + HC-03: P=0.0276). **c)** Images showing PI fluorescence in pericytes immunolabelled for Pdgr-β in control, H_2_O_2_, and H_2_O_2_ plus A96 pre-treatment. Pericyte PI uptake and membrane blebbing are substantially reduced with A96 treatment. Arrow notes a membrane bleb in H_2_O_2_ conditions. Pericytes were identified as Pdgfr-β expressing cells on branch orders ≥1°. **d)** Summary data showing that A96 or HC-03 significantly reduce the abundance of PI labelled pericytes in H_2_O_2_ or diamide (control: *n*=15, *N*=5; H_2_O_2_: *n*=15, *N*=5; H_2_O_2_ + A96: *n=*15, *N*=5; diamide: *n*=15, *N*=5; diamide + A96: *n*=15, *N*=5; diamide + HC-03: *n=*12, *N*=4; Kruskal-Wallis test, KW=60.68, P<0.0001; Dunn’s multiple comparisons test, control vs H_2_O_2_: P<0.0001; H_2_O_2_ vs H_2_O_2_ + A96: P< 0.0001; H_2_O_2_ vs H_2_O_2_ + HC-03: P=0.0084; control vs diamide: P<0.0001; diamide vs diamide + A96: P=0.0005; diamide vs diamide + HC-03: P=0.0106). **e)** *Left*, A PI labelled pericyte showing a prominent membrane bleb protruding from the cell body. The maximum displacement of the bleb from the cell body (dashed line) was used to quantify bleb size. *Middle*, Summary data of bleb size, acquired from all blebbed cells observed across H_2_O_2_ treated brain slices (*n*=63 blebs, *n*=38 cells, *n=* 16 slices, *N*=5 animals). Violin plot summary statistic lines represent the first quartile, median, and 3^rd^ quartile. *Right*, Relative to control, the abundance of blebbed pericytes is significantly increased in H_2_O_2_. Pre-treatment with A96 significantly reduced the abundance of blebbed pericytes in H_2_O_2_ (control: *n*=15, *N*=5; H_2_O_2_: *n*=15, *N*=5; H_2_O_2_ + A96: *n=*14, *N*=5; Kruskal-Wallis test, KW=17.59, P= 0.0002; Dunn’s multiple comparisons test, control vs H_2_O_2_: P= 0.0002; H_2_O_2_ vs H_2_O_2_ + A96: P= 0.0062; control vs H_2_O_2_ + A96: P> 0.9999).

Our data suggests that TRPA1 has a substantial effect on pericyte damage during oxidative stress, which could be explained by a close association between TRPA1 and capillaries. We examined this possibility by performing immunostaining for TRPA1 in glyoxal fixed cortical brain tissue. Imaging of cortical capillaries revealed dense immunolabelling along the length of brain capillaries (Figure 7a,b). This TRPA1 immunolabelling was closely associated with pericyte processes, endothelial cells, and astrocyte endfeet (Figure 7a,b). While TRPA1 has been shown to be have functions in endothelial cells and astrocytes, the presence of the channel in pericytes has not been previously described. To address whether TRPA1 is also present in pericytes, we performed western blotting of lysate from isolated primary rat and human pericytes *in vitro*. Western blotting of cell lysates from human and rat pericytes revealed strong protein bands ranging from ∼110-130 kDa using a TRPA1 antibody (Figure 7c). This molecular weight range is consistent with past studies on TRPA1 ^63^. Blotting was also performed on lysate from primary rat astrocytes, which showed protein bands at the correct molecular weight range, but with a lower abundance than in pericytes (Figure 7c). This data provides evidence that TRPA1 channels are closely associated with brain capillaries and are expressed by pericytes.

**Figure 7.**
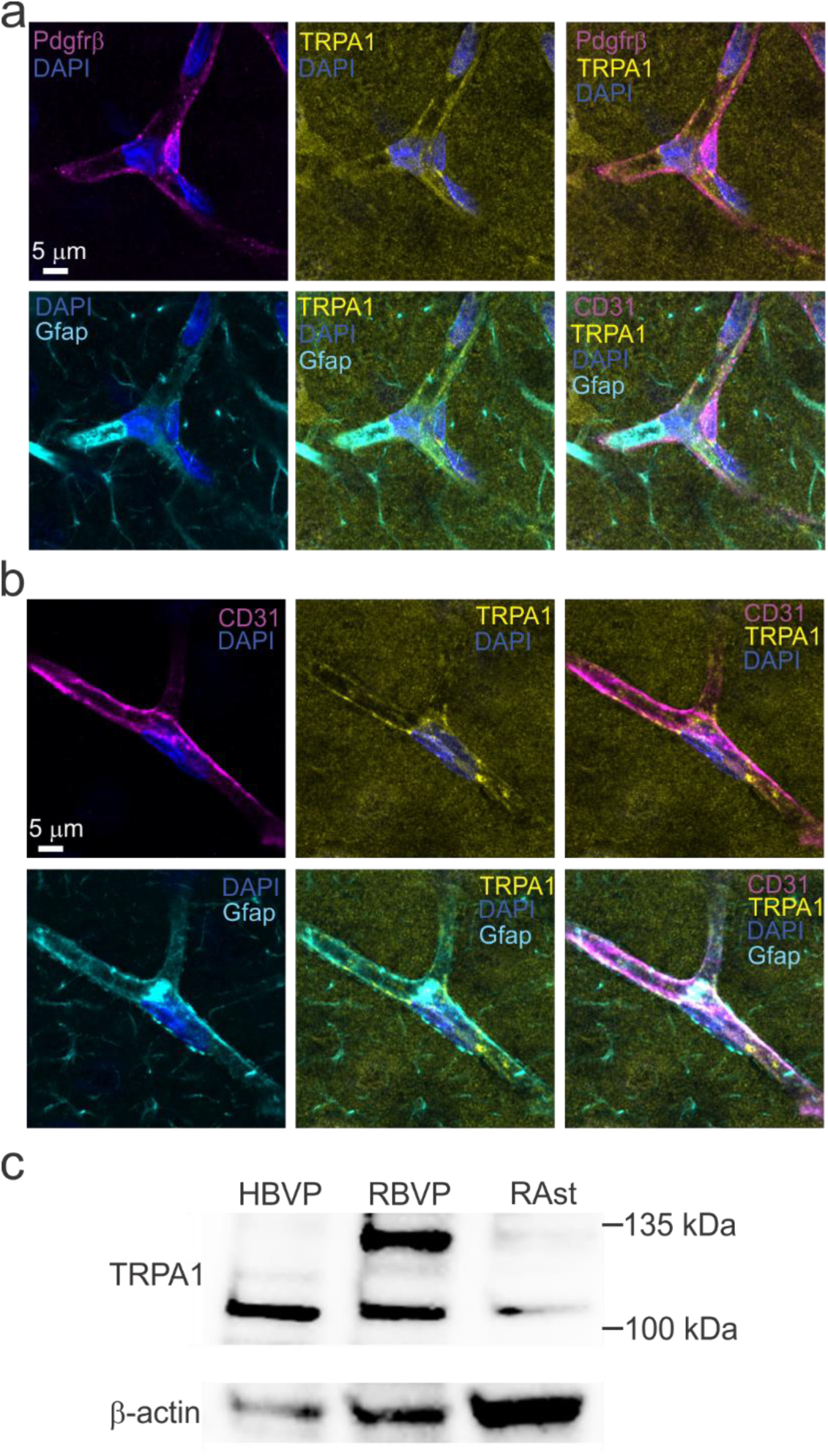
TRPA1 is associated with brain capillaries and is expressed in pericytes. **a)** Representative images of GAA-fixed cortical brain tissue immunolabelled for TRPA1, pericytes with Pdgfr-β, and astrocytes with Gfap. TRPA1 immunolabelling is present along the length of brain capillaries, in close association with Pdgfr-β positive pericyte processes and astrocyte endfeet. **b)** Representative confocal images show the close association of TRPA1 immunolabelling with the endothelial marker CD31 and Gfap labelled astrocyte endfeet. **c)** Western blot showing TRPA1 expression in whole-cell lysates from primary human brain vascular pericytes (HBVP), rat brain vascular pericytes (RBVP), and primary rat astrocytes. Strong immunoreactive bands for TRPA1 are detected at the expected molecular weight (∼110– 130 kDa) in both HBVP and RBVP, with a band of lower intensity observed in astrocytes. β-actin was used as a loading control (bottom panel). Blots are representative of n=7 biological replicates.

Much of our work has utilized PI uptake to assess cell damage in this study, as this is a well-established marker of cell membrane breakdown and death. However, pericytes are known to sequester large molecular weight organic compounds even under physiological conditions. For example, pericytes, but not other cells, are known to sequester the fluorescent molecule NeuroTrace, a derivative of FluoroNissl Green, under physiological conditions via an unknown transport mechanism ^34^. This suggests that the observed PI uptake could be also be driven by changes in the activity of a transport mechanism, rather than simply generalized membrane breakdown. We addressed this possibility by examining whether TRPA1 activity impacts the transport of a large molecular weight compound by pericytes under physiological conditions.

This was tested by examining the impact of TRPA1 antagonists on the uptake of NeuroTrace by pericytes under basal conditions, in the absence of oxidative stress. Brain slices incubated in aCSF in the absence of blockers showed intense labelling of pericyte cell bodies and processes throughout the cortical tissue. In contrast, incubating brain slices with TRPA1 blockers during dye incubation, substantially reduced NeuroTrace labelling of pericytes (Figure 8a). Relative to control conditions, amiloride (1 mM), A96 (20 µM), or HC-03 (10 µM) all significantly reduced the NeuroTrace labelling of pericytes (Figure 8b).

**Figure 8.**
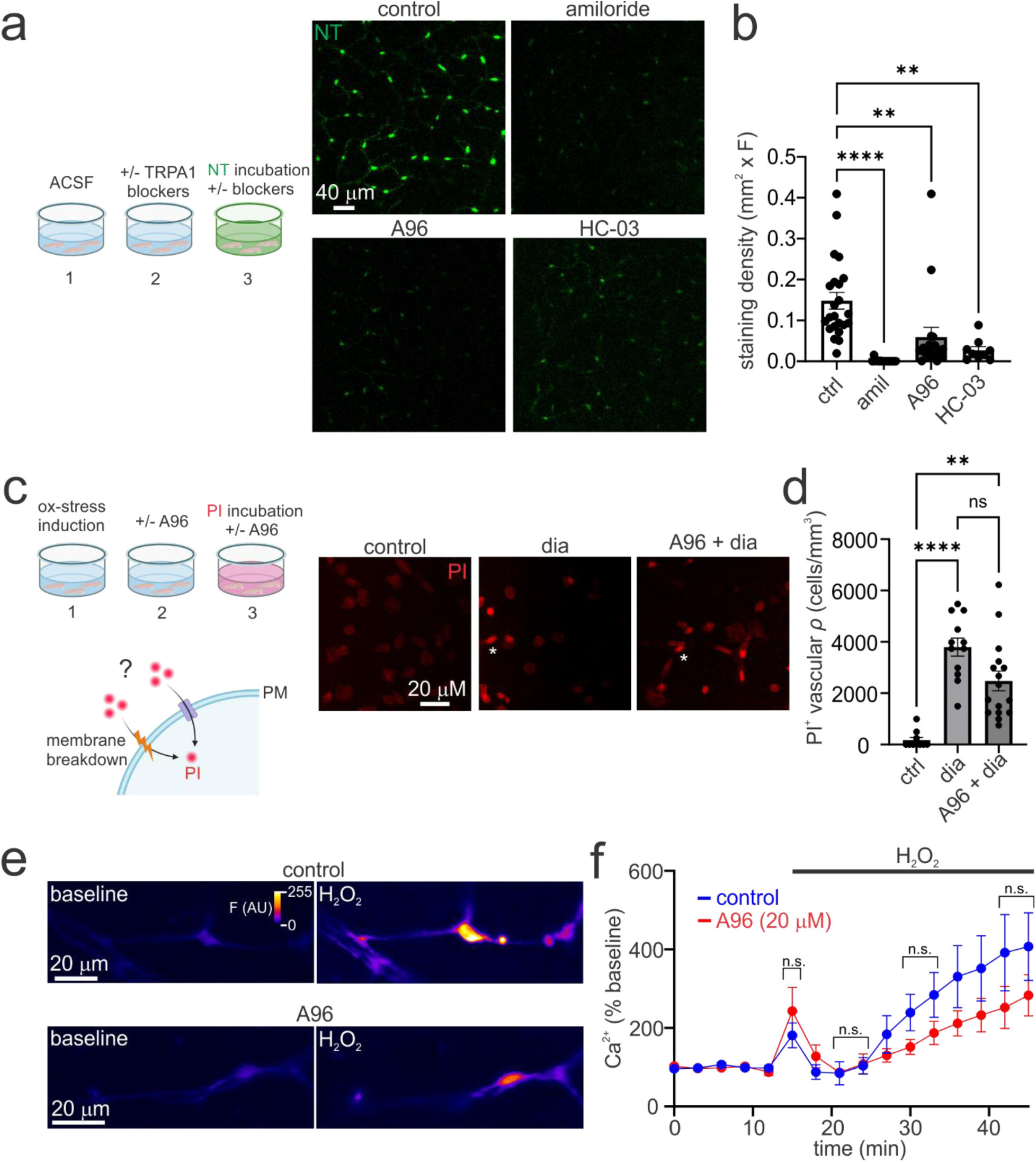
TRPA1 regulates large molecule uptake into pericytes but is not the primary mediator of bulk pericyte Ca^2+^ elevation during oxidative stress. a) *Left*, Acute cortical brain slices were pre-incubated in ACSF or ACSF plus TRPA1 blockers for 20 minutes (**1 & 2**). NeuroTrace (NT) was then added to the bathing solution for 40 min to allow for dye uptake (**3**), followed by washing in ACSF and live two-photon imaging. *Right*, Images depict maximum intensity projections of pericyte NeuroTrace labelling following incubations in control and the following TRPA1 blockers: amiloride (1 mM), A96 (20 µM), and HC-03 (10 µM). Blocking TRPA1 channels during NeuroTrace incubation substantially reduces the intensity of NeuroTrace pericyte labelling. **b)** Summary data showing that incubation in amiloride, A96, and HC-03 significantly reduce NT labelling relative to controls (control, *n*=23, *N*=7; amiloride, *n*=15, *N*=5; A96, *n*=18, *N*=5; HC-03, *n*=9, *N*=3; Kruskal-Wallis test, df= 3, *H*= 46.58, P<0.0001; Dunn’s Multiple comparisons test: control vs amiloride, P<0.0001; control vs A96, P=0.0056; control vs HC-03, P=0.0045). c) *Left*, Brain slices were incubated with diamide (2 mM) to induce oxidative stress for 1 hour. Slices were then incubated for 20 min in control or A96 (20 µM) conditions, followed by the addition of PI to label damaged cells. Lower illustration shows hypothesized routes of PI uptake following oxidative stress induction. *Right*, TRPA1 blockade following oxidative stress induction with diamide does not substantially reduce the number of PI labelled microvascular cells. **d)** Summary data showing that TRPA1 blockade after induction of oxidative stress with diamide does not significantly reduce the abundance of damaged microvascular cells (control, *n*=10, *N*=3; diamide, *n*=12, *N*=3; A96 + diamide, *n*=16, *N*=3; Kruskal-Wallis test, df= 2, *H*= 46.58, P<0.0001; Dunn’s Multiple comparisons test: control vs diamide, P<0.0001; diamide vs A96 + diamide, P=0.1853; control vs A96 + diamide, P=0.0017). **e)** Images depict pericyte GCaMP6f fluorescence in acute brain slices prepared from GCaMP6f; ATP13a5Cre mice. Induction of oxidative stress with H_2_O_2_ (1 mM) produces a short lived Ca^2+^ transient followed by a protracted Ca^2+^ rise in the cell body and processes, which is largely undisturbed by ongoing TRPA1 blockade with A96 (20 µM) **f)** Summary data of pericyte somatic Ca^2+^ levels during H_2_O_2_ application. Incubating brain slices with A96 to block TRPA1 channels, partially reduces the protracted Ca^2+^ response in H_2_O_2_, but this difference does not reach statistical significance at any time point (H_2_O_2_: *n*=7, *N*=7; H_2_O_2_ + A96: *n*=5, *N*=5; unpaired student’s t tests).

Our results indicate that TRPA1 can regulate the uptake of large molecules by pericytes through a transport mechanism. Thus, we next tested whether the PI uptake into pericytes during oxidative stress could also be mediated by the TRPA1-dependent transport system. In our earlier experiments, brain slices were pre-treated with A96 prior to oxidative stress induction and PI loading. Consequently, it is difficult to discern whether the reduced PI labelling occurred because A96 directly blocked TRPA1-dependent dye transport or because it disrupted a protracted TRPA1-dependent signaling cascade that leads to secondary membrane breakdown and PI uptake. If TRPA1-dependent transport is the primary mechanism underlying PI uptake into pericytes, post-oxidative stress TRPA1 blockade should still effectively prevent dye uptake. To test this, brain slices were first exposed to diamide for 1 hour in dye-free aCSF to establish oxidative stress (Figure 8c). Slices were then incubated with or without A96, followed by incubation in PI, tissue fixation and imaging (Figure 8c). As seen previously, oxidative stress alone produced an increase in PI-labelled microvascular cells (Figure 8c,d). However, applying A96 after oxidative stress induction caused only a partial, non-significant reduction in the density of PI-positive cells (Figure 8c,d). These data indicate that PI uptake during acute oxidative stress is not primarily mediated by the TRPA1 dependent transport mechanism, but instead reflects a protracted downstream mechanism, such as generalized membrane breakdown.

Pericyte damage during oxidative stress required TRPA1 activity and Ca^2+^ influx, suggesting that TRPA1 may initiate Ca^2+^ responses in pericytes during oxidative stress. We therefore examined evidence for a TRPA1-dependent Ca^2+^ response to oxidative stress in these cells. To do this, we prepared acute brain slices from GCaMP6f; ATP13a5Cre mice. These mice express Cre-recombinase specifically in brain pericytes, leading to pericyte specific GCaMP expression ^64^. Application of H_2_O_2_ produced a transient Ca^2+^ response in the soma and processes followed by a protracted Ca^2+^ elevation which progressed over many minutes (Figure 8e,f).

Incubating brain slices with A96 (20 µM) during H_2_O_2_ application revealed a slight, but not significantly reduced, Ca^2+^ response (Figure 8e,f). This data suggests that TRPA1 channels are not the primary mechanism underlying ROS-induced global Ca^2+^ elevation in pericytes, despite playing a major role in cell damage.

## Discussion

Identifying the cellular and molecular mechanisms that protect against oxidative stress is critical to developing targeted therapeutic strategies for neurological disease. In this study we demonstrate that pericytes exhibit marked sensitivity to oxidative stress. Examination of the mechanisms involved in pericyte injury showed that this process is non-apoptotic, non-necroptotic, and dependent on Ca^2+^ influx. Through pharmacological analyses, we establish that TRPA1 channels serve as essential mediators of oxidative damage to pericytes (Figure 9). We further show that TRPA1 regulates basal large-molecule uptake by pericytes, but is not the primary driver of ROS evoked bulk cytosolic Ca^2+^ pericytes. Collectively, this work highlights pericytes as a critical cell type disrupted by oxidative stress in the brain and suggests TRPA1 antagonism as a promising therapeutic avenue to preserve pericyte and microvascular integrity in acute and chronic brain disorders.

**Figure 9.**
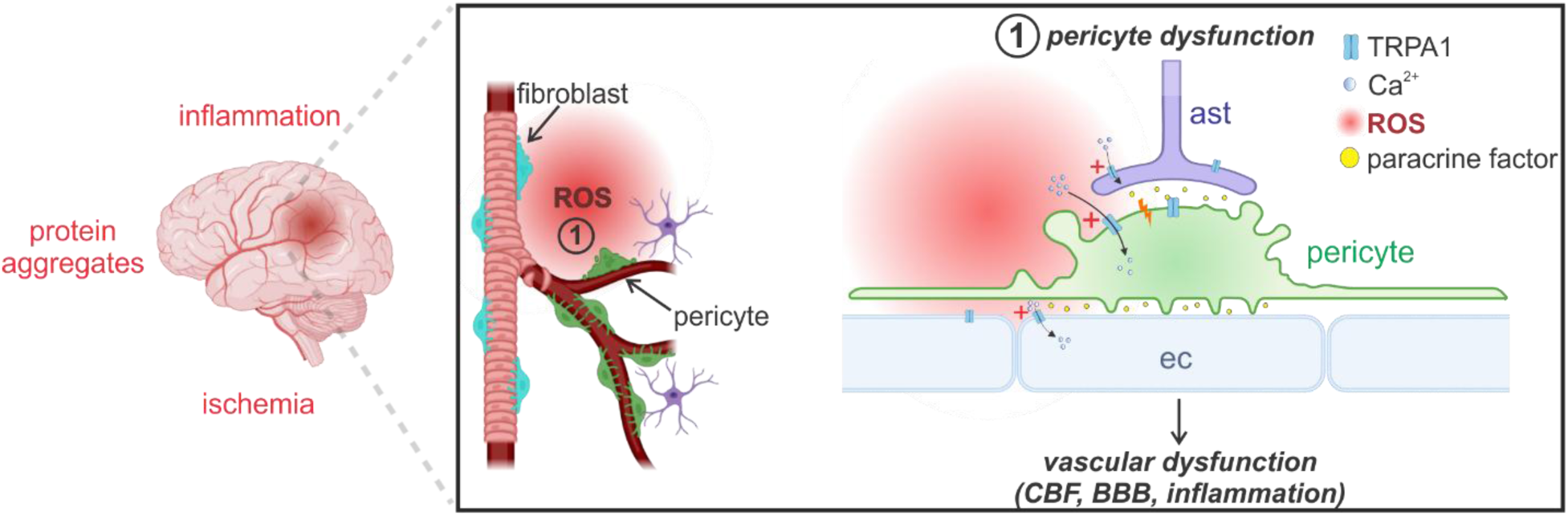
Model of TRPA1-dependent pericyte damage during oxidative stress. Visual abstract depicting a proposed mechanism of oxidative stress impacts during brain pathology in accordance with the presented results. Pericytes and perivascular fibroblasts are sensitive, early responders to pathological ROS elevation in the brain. ROS leads to the activation of TRPA1 channels, either in pericytes or neighboring endothelial cells and astrocytes, leading to Ca^2+^ influx. The resultant pericyte cell blebbing and membrane damage occurs either through direct TRPA1 activity in pericytes or a TRPA1-dependent release of paracrine factors from astrocytes and endothelial cells. By disrupting pericyte integrity, the ROS-TRPA1 axis could contribute to pathological changes in cerebral blood flow, BBB, and vascular inflammation. Image produced using BioRender.com.

### Pericytes and perivascular fibroblasts are damaged during oxidative stress

Our results demonstrate that acute oxidative stress triggers substantial PI uptake in both pericytes and perivascular fibroblasts. Cellular uptake of PI is a well-established marker of plasma membrane compromise and a reliable predictor of cell death ^14,24^. For example, following experimental traumatic brain injury, PI-positive cells exhibit structural damage and are eventually cleared from brain tissue ^24^. The robust PI uptake observed in pericytes and perivascular fibroblasts during oxidative stress indicates a state of severe cellular stress and irreversible membrane breakdown. Consistent with this, damaged pericytes displayed pronounced membrane blebbing protruding away from the vessel wall into the parenchymal space, likely reflecting herniation through the vascular basement membrane. Similar pericyte blebbing has been observed *in vivo* during pericyte pathology induced by ischemic stroke and cellular overactivation ^33,65^. While we discovered that TRPA1 regulates the transport of the large compound, NeuroTrace, into pericytes under physiological conditions, our data suggests that the observed PI uptake during oxidative stress is largely independent of such transport activity.

Applying TRPA1 blockers after establishing oxidative stress produced only a minor, non-significant reduction in PI uptake. This indicates that PI uptake into pericytes requires TRPA1 activation, but occurs through a secondary, downstream mechanism rather than a regulated transport process directly coupled to channel activity. Based on the severe membrane distortion observed in pericytes, membrane lysis is the likely cause of PI uptake observed in our study.

However, a contribution from secondary large-pore pathways, such as pannexins ^66^ cannot be excluded. Regardless of the exact mechanism, these data provide clear evidence that the observed PI labelled cells enter a damaged state during acute oxidative stress. Furthermore, these findings highlight the need for careful interpretation when using fluorescent dye uptake assays to assess pericyte ROS generation, viability, and death, as has been done previously ^42,67,68^.

### Pericytes and perivascular fibroblasts exhibit heightened vulnerability to oxidative injury

Our findings indicate that pericytes and perivascular fibroblasts possess an intrinsic sensitivity to oxidative damage compared to neighboring cells. An initial indicator of this vulnerability was that damaged microvascular cells, corresponding primarily to Pdgfr-β positive cells, accounted for a high proportion of all damaged cells in H_2_O_2_-treated cortical tissue, and comprised the majority of damaged cells following diamide exposure. This disproportionate representation was unexpected, as vascular cells constitute only a minor fraction of the total cell population in cortical layers I and II (∼32% in layer I and ∼12% in layer II ^30^). Thus, the high density of damaged microvascular cells is likely not accounted for by regional differences in cell density alone, pointing instead to heightened susceptibility to oxidative stress. Comparisons within specific vascular compartments provide more evidence for this heightened vulnerability. Along vessel branches of order ≥ 1° (ACT, mid-capillary bed, and post-capillary venules), damaged pericytes substantially outnumbered damaged endothelial cells. Given that the pericyte-to-endothelial cell ratio in the rodent cortex ranges from 1:1 to 1:4 ^69,70^, this disparity cannot be attributed to differences in cell abundance and likely reflects comparatively high sensitivity of pericytes to oxidative stress. Similarly, on primary-order penetrating vessels (0°), diamide induced robust PI uptake in perivascular fibroblasts while leaving adjacent smooth muscle cells largely intact. Importantly, we cannot rule out the possibility that other cells, such as endothelial cells or smooth muscle cells, are undergoing non-necrotic forms of injury and cell death in our model. For example, non-lytic modes of cell death or stress do not necessarily involve membrane permeabilization ^71,72^. Thus, our data provide evidence that capillary pericytes and perivascular fibroblasts represent microvascular cell types which are more prone to undergoing acute, oxidative stress-induced necrotic phenotype than other cells.

Differential vulnerability to oxidative stress is a previously recognized phenomenon among discrete cell populations in the central nervous system. For example, dopaminergic neurons of the substantia nigra pars compacta exhibit greater susceptibility to oxidative stress and degeneration than adjacent dopaminergic neurons in the ventral tegmental area ^19,73^. This selective sensitivity is thought to arise in part from elevated basal oxidative stress driven by sustained Ca^2+^ influx^19,73^. Similarly, among glial populations, oligodendrocytes display marked sensitivity to oxidative damage compared to more resilient astrocytes ^6,74,75^. These glial differences have been attributed to a comparatively lower reduced glutathione pool and higher iron content in oligodendrocytes compared to astrocytes ^75,76^. Our findings indicate that oxidative stress sensitivity in pericytes can be accounted for by close functional coupling between TRPA1, an intrinsically ROS-sensitive non-selective cation channel, and pericytes. We provided evidence that TRPA1 is closely associated with pericytes on brain capillaries *in vivo* and is expressed in isolated primary brain pericytes *in vitro*. Moreover, pharmacological blockade of TRPA1 markedly attenuated the basal uptake of NeuroTrace into pericytes, consistent with a high functional relevance of TRPA1 activity in these cells. It is also possible that the heightened vulnerability of pericytes and fibroblasts observed here may be impacted by intrinsic differences in antioxidant buffering capacity, distinct enzymatic ROS-generating systems, elevated baseline metabolic stress, or an abundance of downstream redox-sensitive targets that trigger cytotoxic cascades. There is little known about redox homeostasis in pericytes and perivascular fibroblasts in the brain. Notably, previous studies have demonstrated that amyloid-β oligomers evoke substantially greater ROS accumulation in capillary pericytes than in adjacent parenchymal cells ^68^. While the exact biochemical drivers of this differential response remain unresolved, they could stem from distinct rates of ROS production or clearance within pericytes.

### Pericyte dysfunction during oxidative stress is driven by TRPA1 ion channels and Ca^2+^

Previous work has demonstrated an impact of various Ca^2+^ entry pathways on pericytes in health and disease, including voltage-gated Ca^2+^ channels, TRPC channels, and ORAI channels ^51,67,77,78^. However, we found that blocking voltage-gated Ca^2+^ channels, TRPC/ORAI channels, TRPV2, and intracellular Ca^2+^ release had no protective effect against pericyte oxidative injury. In contrast, pericyte damage was significantly reduced by ruthenium red, amiloride, benzamil, HC-03, and A96. This pharmacological profile strongly points to an involvement of TRPA1 channels in pericyte damage ^57,79^. While ruthenium red and amiloride derivatives can engage other channel families at higher concentrations, HC-03 and A96 are well-established TRPA1 antagonists ^62^. A96 in particular exhibits high selectivity for TRPA1 over other TRP channel subtypes and non-TRP targets ^62^. Although definitive validation will require cell-specific gene knockdown or knockout models in future studies, our pharmacological data robustly implicate TRPA1 in pericyte damage during oxidative stress.

The molecular cascade coupling ROS, TRPA1 activation, Ca^2+^ influx, and pericyte damage is unclear. A simple model to explain our findings is that ROS activates TRPA1 in pericytes via covalent modification of intracellular cysteines on the channel ^55^, leading Ca^2+^ influx, pericyte blebbing, and membrane disruption (Figure 9). Consistent with this model, we provided evidence for TRPA1 expression in brain pericytes via western blotting of cultured pericyte lysate. However, examination of intracellular Ca^2+^ in pericytes during TRPA1 antagonism with A96 showed that there was only a slight, non-significant reduction in H_2_O_2_-evoked bulk Ca^2+^ elevations. This demonstrates that, despite underlying pericyte damage, TRPA1 is not the primary driver of global intracellular Ca^2+^ increases during oxidative stress. Other pathways must account for this bulk cytosolic Ca^2+^ elevation under these conditions; for example, H_2_O_2_ has been shown to evoke pericyte Ca^2+^ rises and constriction via voltage-gated Ca^2+^ channels and downstream endothelin-1A receptor signaling, respectively ^77^. If TRPA1-dependent toxicity occurs in a pericyte autonomous manner, one explanation for this apparent discrepancy is that TRPA1 signaling is triggered by spatially confined Ca^2+^ microdomains, rather than bulk cytosolic accumulation. Such close physical coupling between Ca^2+^ entry and downstream effectors is a well-described property of intracellular Ca^2+^ signaling ^80^. Notably, astrocytic TRPA1 channels generate dynamic Ca^2+^ microdomains that are readily detected by membrane-tethered GCaMP constructs, but are not readily measured when using cytosolic GCaMP reporters ^60,61,81^. Similarly, our use of a cytosolic GCaMP6f sensor may have been insufficient to resolve spatially restricted TRPA1-mediated Ca^2+^ signals. Localized Ca^2+^ influx through TRPA1 could potentially recruit Ca^2+^-dependent enzymes, such as calpains, leading to cytoskeletal breakdown, membrane blebbing, and subsequent lysis ^82,83^. An alternative mechanism by which TRPA1 can promote pericyte damage is through a multi-cellular cascade involving paracrine signals derived from astrocytes and endothelial cells, which express functional TRPA1 channels. In astrocytes, TRPA1 regulates resting Ca^2+^ levels and the release of vasoactive molecules that modulate neighboring cells ^59–61^. Brain endothelial cells also express TRPA1 and are capable of releasing factors that influence pericytes, such as endothelin-1 and matrix metalloproteases ^58,68,84,85^. Further work utilizing membrane-targeted biosensors and cell-type-specific genetic tools will be essential to dissect the coupling between TRPA1 activation and pericyte degeneration.

### Conclusions and Implications

Our results suggest that the ROS-TRPA1 axis could contribute to pericyte dysfunction and death in the numerous brain disorders associated with oxidative stress. This has critical functional implications given the essential roles pericytes play within the cerebral microvasculature. Pericytes actively regulate local capillary diameter and cerebral blood flow through their contractile machinery ^31,42^. Furthermore, there is evidence that pericytes govern the maintenance of the blood-brain barrier during development ^86^, although this role may be less critical in adulthood ^87^. Beyond hemodynamic control, pericytes modulate parenchymal function through paracrine signaling ^88,89^, and serve as a source of mesenchymal stem cells to facilitate brain tissue repair following injury ^35^.

Under pathological conditions, pericytes undergo a number of phenotypic alterations, including hyperconstriction^33,42^, inflammatory cytokine release ^88^, and structural degeneration and death ^39,41^. Notably, in ischemic stroke, Alzheimer’s disease, and diabetic retinopathy, pericytes display marked vulnerability to degeneration and loss while the adjacent endothelium often remains structurally intact ^41,42,90,91^. This selective loss of pericytes precipitates aberrant blood flow distribution, capillary regression, and blood-brain barrier disruption ^87,89^. Protecting pericytes from structural breakdown therefore represents an important therapeutic avenue for preserving microvascular integrity in neurovascular and neurodegenerative disorders. Our findings indicate that the heightened sensitivity of pericytes to oxidative stress is mediated by TRPA1, which may contribute to diverse facets of pericyte pathology, including pathological constriction, inflammatory signaling, and cell loss. Targeting TRPA1 represents a compelling strategy to preserve pericyte viability and maintain capillary patency, particularly during acute reperfusion injury or chronic neurodegeneration, as seen in Alzheimer’s disease.

Finally, our findings suggest that TRPA1 governs the uptake of the selective pericyte marker NeuroTrace. Under physiological conditions, NeuroTrace specifically labels brain pericytes over neighboring endothelial cells, vascular smooth muscle cells, glia, and neurons ^34^. The molecular mechanisms underlying this dye sequestration has not been previously identified, but are presumed to reflect a transport process unique to pericytes. In this study, we demonstrate that multiple distinct TRPA1 antagonists suppress basal NeuroTrace uptake by pericytes, suggesting a critical role for TRPA1. The channel could govern this process by direct passage through the large conductance TRPA1 pore ^92,93^ or by downstream regulation of a transport system in pericytes. As NeuroTrace uptake is highly specific to pericytes, it suggests that elevated TRPA1 activity or a TRPA1-dependent transport system is a feature unique to pericytes. Similar to TRPA1 in peripheral tissue, the strong functional coupling between TRPA1 and pericytes may allow these cells to readily sense and respond to the various noxious activators of TRPA1, including ROS, oxidized lipids, nitric oxide, and the bacterial lipopolysaccharide ^62,79,94–99^ . In this capacity, pericytes could serve as early responders to noxious stimuli originating from the bloodstream, endothelium, or parenchyma. Future investigations utilizing genetic knockdown models will be vital to resolve the functional impacts of TRPA1 on pericytes and fully establish its therapeutic potential in cerebrovascular disease.

## Acknowledgements

This work was supported by grants awarded to B.A.M from the Alzheimer’s Association (VCID-UMD-26-1529075), Canadian Institutes of Health Research (CIHR), and Parkinson Canada.

---

**Figure S1.**
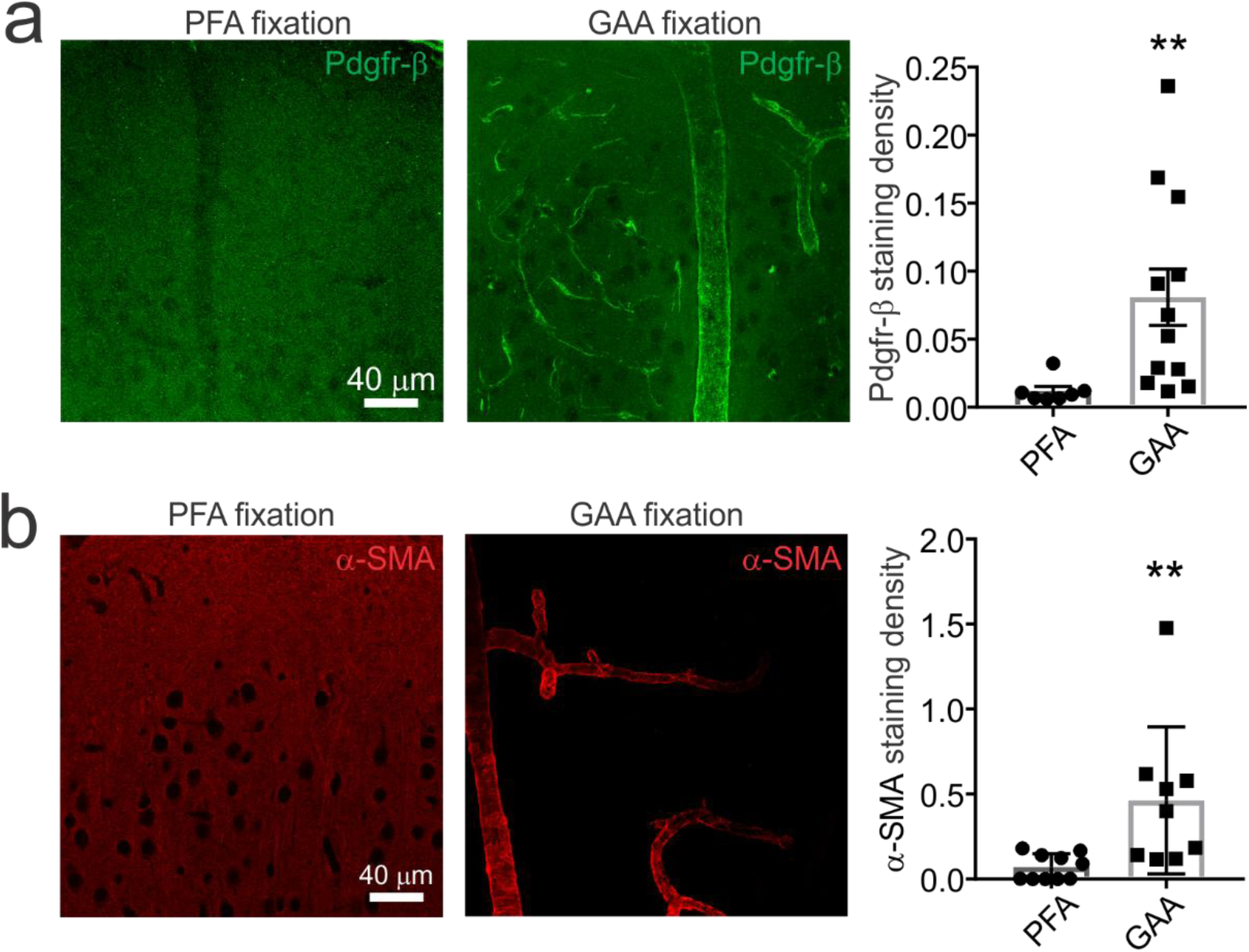
Glyoxal fixation of brain tissue improves mural cell immunolabelling. **a)** *Left*, Images depict maximum intensity projections of Pdgfr-β stained cortical tissue fixed with paraformaldehyde (PFA) or glyoxal, acetic acid (GAA). *Right*, The Pdgfr-β staining density is significantly greater in GAA fixed tissue relative to PFA fixed tissue (PFA: *n*=7, GAA: *n*=12; Mann-Whitney test, *U*=6, P=0.0012). **b)** *Left*, Images depict maximum intensity projections of α-SMA immunostaining in PFA and GAA fixed tissue. *Right*, The staining density of α-SMA is significantly greater in GAA fixed vs PFA fixed tissue (PFA: *n*=12, GAA: *n*=11; Mann-Whitney test, *U*=10, P=0.0002).

